# Zika virus reprograms the host tRNA epitranscriptome to adapt translation to A-ending codon bias

**DOI:** 10.1101/2025.06.03.657606

**Authors:** Patrick Eldin, Eric Bernard, Cathrine B. Vågbø, Sara Salinas, Lynn George, Masahiko Ajiro, Masatoshi Hagiwara, Geir Slupphaug, Laurence Briant

## Abstract

The codon usage bias of the Zika virus (ZIKV) genome is skewed towards AA-ending codons, which are preferentially decoded by U_34_-modified cognate tRNAs. This contrasts with the human host’s preference for AG-ending codons, suggesting that ZIKV may exploit specific tRNA modifications to optimize protein synthesis within human cells. To test this hypothesis, we used codon-biased GFP sensors and found that ZIKV infection transiently increased the expression of AA-biased GFP at the expense of AG-biased GFP. Mass spectrometry analysis further showed that ZIKV virus infection increases mcm^5^U34 and mcm^5^s^2^U_34_ tRNA modification content in host cells. In ELP1-deficient cells, which exhibit reduced U_34_ modifications, ZIKV replication was impaired. Enhancing U_34_ modification through using a small molecule known to restore ELP1 expression, rescued viral replication in these cells. Moreover, CRISPR/Cas9 and shRNA-mediated knockdown of key enzymes involved in U_34_ modification, ELP1, ALKBH8, and CTU1, significantly reduced ZIKV replication. Collectively, these results provide strong evidence that ZIKV reprograms the host tRNA epitranscriptome and exploits host cell tRNA modifications, particularly at the wobble position U_34_, to optimize translation of its own proteins and promote viral replication.

Graphical abstract
Zika virus (ZIKV) reprograms the host cell’s tRNA epitranscriptome by altering U_34_ modifications, specifically enhancing mcm^5^s^2^U levels, thereby adapting the host tRNA pool to match its skewed codon bias and facilitate efficient viral protein synthesis.

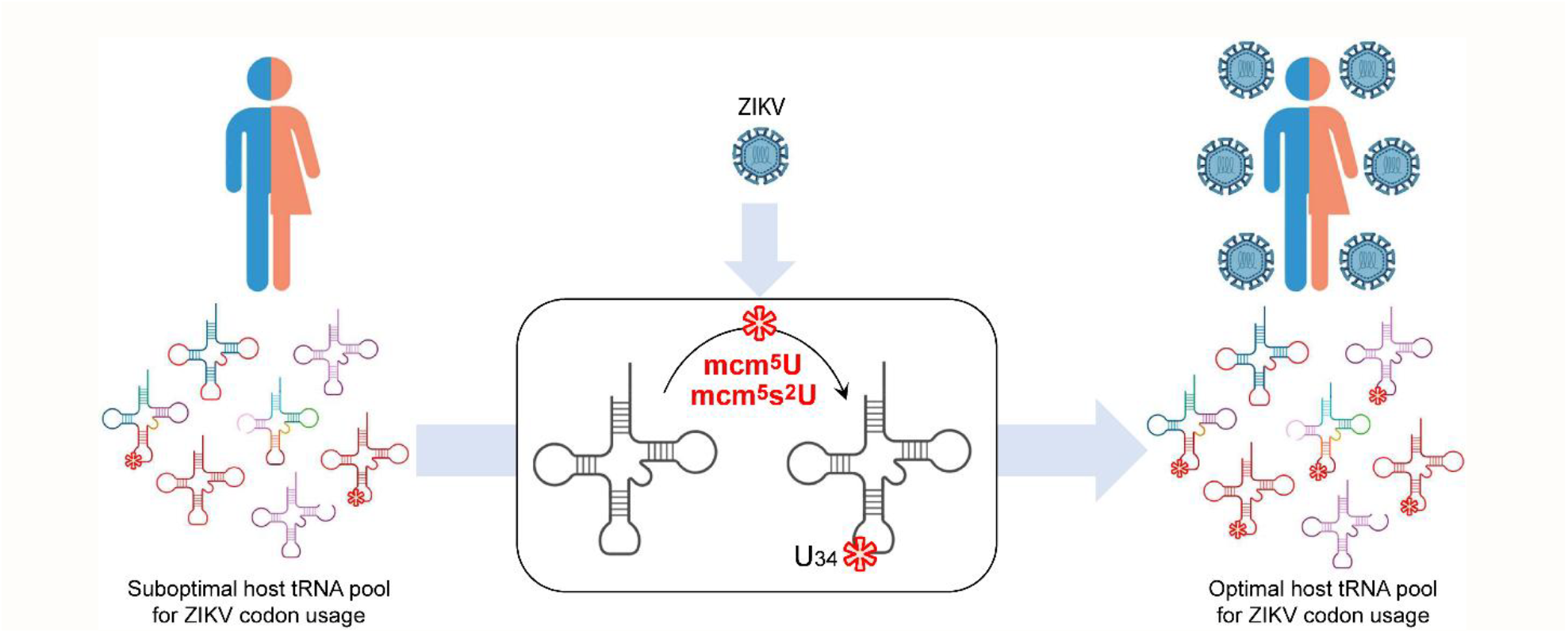

## Introduction

Zika virus (ZIKV), a mosquito-borne *flavivirus*, causes a wide range of neurological complications including fetal microcephaly and Guillain-Barré syndrome. First identified in 1947 in Uganda, ZIKV remained confined to sporadic human cases in Africa and Asia until recently, when large outbreaks successively occurred in the Pacific Yap Islands (2007), French Polynesia (2013) and South America (2015). In 2015, the widespread presence of ZIKV primary *Aedes sp*. vectors across Latin America, combined with favorable climatic conditions, created the ideal environment for a massive outbreak in Brazil, which rapidly spread across Latin America and the Caribbean (1). The global distribution and continued geographic expansion of these mosquito vectors underscore the urgent need for a coordinated response to prevent future ZIKV outbreaks and limit viral spread.

ZIKV is a positive-sense single-stranded RNA virus with a genome of ~10.7 kb. It contains a single long open reading frame encoding a polyprotein (5’-C-prM-E-NS1-NS2A-NS2B-NS3-NS4A-NS4B-NS5-3’), flanked by non-coding regions (5’- and 3’-UTRs). This polyprotein is processed into three structural proteins, capsid (C), precursor of membrane (prM), and envelope (E), and seven non-structural (NS) proteins (2). Upon entry into target cells, ZIKV strictly depends on the host’s translational machinery for the early synthesis of ZIKV NS proteins required for replication and, subsequently, for the production of new viral progeny (3). In this process, tRNAs are key effector molecules bridging the genetic code and amino acid identity. Since the genetic code is highly degenerate, the usage of synonymous codons, referred to as codon usage bias (CUB), can greatly differ between cell types or organisms. This bias is echoed by the composition and abundance of the cellular tRNA pool. Many viruses, including RNA viruses, display a codon bias incompatible with the tRNA pool of their host cells, posing a potential barrier to efficient viral protein synthesis and consequently to their propagation (4, 5). To overcome this major obstacle, viruses may either adapt their codon usage to match that of their hosts (6, 7)or remodel the host tRNA pool to favor their own codon preferences (8–10).

The spectacular emergence of Asian-genotype ZIKV strains during recent outbreaks may at least in part rely on the evolutionary optimization of this vital translation step aimed at optimizing viral translation both in their human host and mosquito vector (11). Nevertheless, analysis of codon-based evolution using eleven ZIKV strains from different geographical locations has revealed only a modest overall codon usage bias, with a consistent preference for A/U-ending codons (12, 13), which contrasts with the human genome’s preference toward C/G-ending codons (14). This mismatch poses a potential challenge for efficient viral translation, especially when host tRNAs corresponding to these preferred viral codons are underrepresented or lack necessary modifications. Since the efficient decoding of 11 out of the 14 A-ending codons used in humans require post-transcriptional modifications (PTMs) at the wobble U_34_ position of the cognate tRNAs (15) (Figure 1A), the proper modification of these U_34_-tRNAs by the adequate cellular enzymatic machinery may be an important factor influencing the efficiency of ZIKV genome translation in human cells.

**Figure 1.**
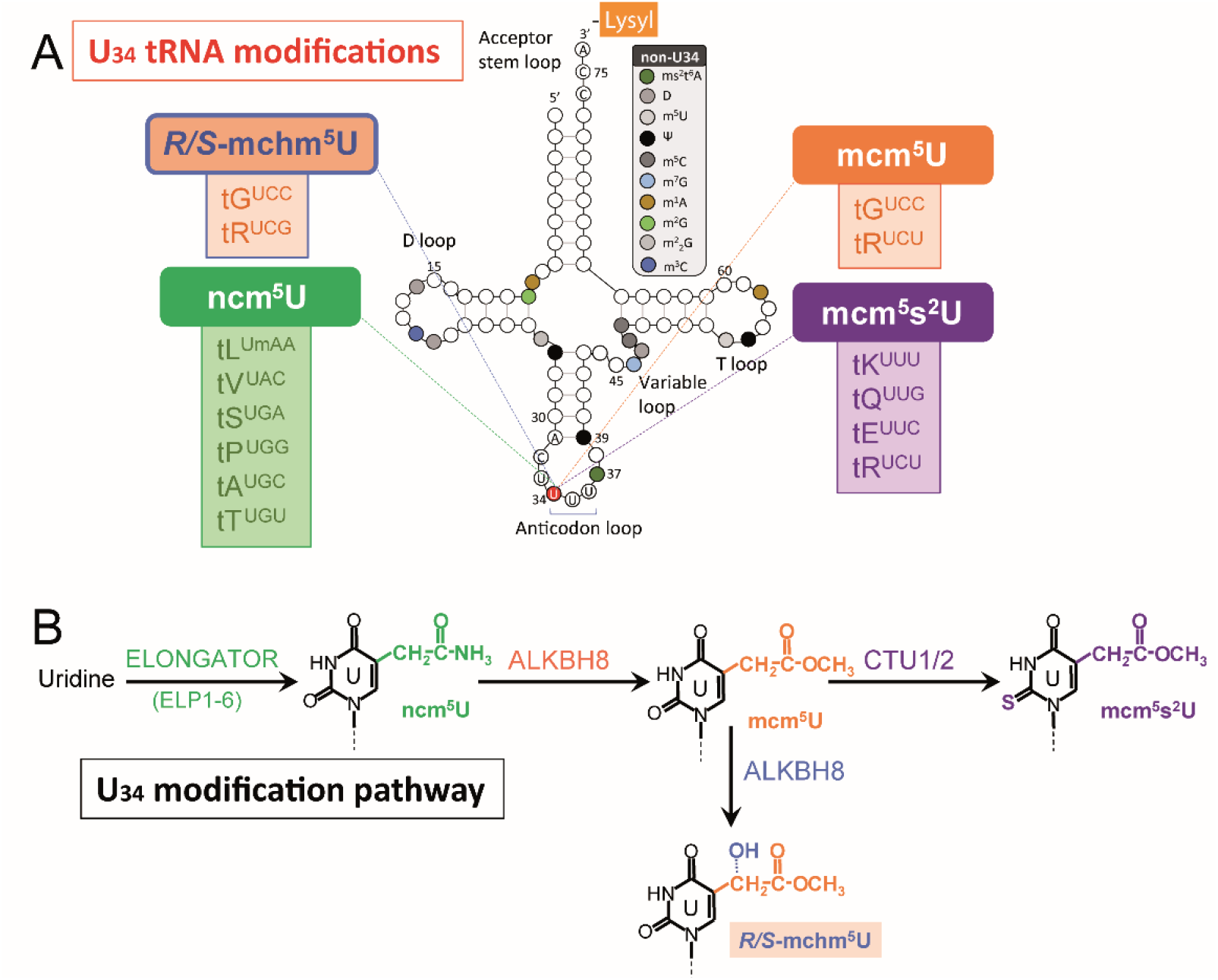
U_34_ tRNA modifications required for decoding AA-ending, U_34_-sensitive codons for Lys, Gln, Glu and Arg. (A) Modifications at the wobble uridine (U_34_) position of tRNA molecules are highlighted with colored boxes corresponding to each modification. Modifications occurring outside U_34_ are boxed in grey. (B) Enzymatic pathway responsible for U_34_ tRNA modifications. A subsequent hydroxylation of mcm^5^U by the oxygenase activity of ALKBH8 can generate mchm^5^U *R/S*-diastereoisomers found on tG^*UCC*^ and tR^*UCG*^. Note that tR^*UCU*^ can carry either mcm^5^U or mcm^5^s^2^U modification.

This machinery (Figure 1B) involves the coordinated activities of the conserved six-subunit acetyltransferase ELONGATOR protein complex (ELP1-6), the methyltransferase ALKBH8 and the thiouridylase complex CTU1–CTU2. Together with the ubiquitin-related modifier (URM) pathway (16–18), these enzymes catalyze the addition of 5-carbamoylmethyl-(ncm^5^), 5-methoxycarbonylmethyl (mcm^5^), a 2-thiol (s^2^), or both (mcm^5^s^2^) at the wobble U_34_ position of tRNAs. These PTMs are critical to enhance codon recognition and decoding efficiency, thereby enabling efficient translation of A-ending codons that otherwise impose suboptimal codon– anticodon interactions (19–23). It is typically the case for codons encoding lysine (AAA/G), glutamine (CAA/G), and glutamic acid (GAA/G), each encoded by two synonymous codons ending in either -AA or -AG. While AG-ending codons are decoded by unmodified cognate tRNAs (tK^*CUU*^, tQ^*CUG*^ and tE^*CUC*^, respectively), AA-ending codons are decoded by tRNA species with a U at the wobble position (U_34_) (tK^*UUU*^ for lysine, tQ^*UUG*^ for glutamine, tE^*UUC*^ for glutamic acid). Although these tRNAs can recognize both the AA- and AG-ending versions of these codons, efficient decoding of the AA-ending triplet strictly requires the presence of the mcm^5^s^2^ modification at U_34_ (24, 25). While the mcm^5^s^2^ modification refines U_34_ recognition and specifically enhances the translational efficiency of problematic AA-ending codons that impose restrictive codon– anticodon interactions (25), its effect on AG-ending codons is debated, with some studies suggesting no impact and others indicating a decrease in translational efficiency (26–29). In addition to modifying tK^*UUU*^, tQ^*UUG*^, and tE^*UUC*^ in humans, the mcm^5^s^2^U_34_ modification is also found on the tR^*UCU*^ specific for arginine, where it is essential for the efficient decoding of the Arg^*AGA*^ codon (30–34).

Any disparity between ZIKV codon usage and the composition or modification state of the host tRNA pool, particularly inadequate levels of U_34_ modifications, could result in stalled translation, mistranslation, and proteotoxic stress, which would ultimately be detrimental to the virus replication. Idiosyncratic recognition of other U_34_-sensitive AA-ending codons, including Arg (AGA/G), Ser (UCA/G) or Pro (CCA/G), by their cognate tRNAs (tR^*UCU*^ and tS^*UGA*^ or tP^*UGG*^, respectively), also requires proper U_34_ modification to establish stable contact with their cognate A-ending codons (35). In the absence of these modifications, the tRNAs cannot efficiently induce the expected conformational changes in the small ribosomal subunit (SSU), disrupting translation elongation, protein synthesis and potentially hindering viral propagation (36).

In this study, we investigated whether ZIKV manipulates the host’s tRNA modification machinery to optimize the translation of its A-ending codon-biased genome. We first reassessed codon usage patterns across different ZIKV genotypes (African and Asian) and compared them with those in the human host, paying particular attention to codons recognized by U_34_-modified cognate tRNAs. This analysis suggests that optimal ZIKV translation in human cells requires adjustments of the host tRNA pool, either through the upregulation of U_34_-containing tRNAs or through increased U_34_ modification within U_34_-containing tRNA subtypes. To functionally assess this hypothesis, we employed codon-biased GFP reporter constructs delivered *via* VLP transduction or mRNA transfection and demonstrated that ZIKV infection transiently enhanced the expression of an AA-biased GFP construct relative to an AG-biased counterpart. Consistently, mass spectrometry analysis of modifications in the small RNA fraction of ZIKV-infected cells confirmed an increased abundance of mcm^5^s^2^U_34_ modification. The absolute necessity for ZIKV to optimize decoding of its A-ending codon-biased genome through U_34_ modification of host tRNA pool was clearly demonstrated in ELP1-deficient human primary cells. Here, the intrinsic hypomethylation of U_34_-tRNAs significantly impaired ZIKV translation and spread. Strikingly, treatment with RECTAS (<u>RECT</u>ifier of <u>A</u>berrant <u>S</u>plicing), a small molecule which restores ELP1 expression and normalizes U_34_ tRNA modification levels in ELP1-deficient human primary cells, rescued the cell’s permissiveness to ZIKV infection. Concurrently, targeted disruption of each of the three key enzymes in U_34_ modification, ELP1, ALKBH8 and CTU1, using CRISPR/Cas9 and shRNA approaches, further underscored the virus’s strong dependency on a properly modified tRNA pool for efficient translation. Collectively, these findings highlight the essential role of U_34_ modification containing tRNAs in the efficient decoding of ZIKV’s codon-biased genome and establish a direct functional link between the host tRNA epitranscriptomic regulation and viral replication efficiency.

## Material and Methods

### Viruses and cells

PF13, and MR766 ZIKV molecular clones were obtained from Mathew Evans (37). BeH8 (mCherry reporter) ZIKV replicon (38) was a generous gift of Andres Merits (University of Tartu, Estonia). Virus production and titration were performed in Vero cells. HeLa, A549, HEK293T, Vero, SH-SY5Y cells and human primary astrocytes (obtained from ScienCell) were cultured in DMEM Glutamax medium (GIBCO) supplemented with Penicillin/Streptomycin and 5 % fetal calf serum. Patient primary fibroblasts were from Coriell Institute: GMO1652 was derived from non-FD control [Skin fibroblast (arm) from 11 years old Caucasian female]; GMO4959 derived from FD patient [Skin fibroblast (arm) from 10 years old Caucasian female]. ZIKV strains are displayed in Supplementary Table 1. Infections were done at the indicated MOIs and checked routinely by RTqPCR as outlined in Supplementary Figure S5.

To monitor ZIKV translation, we used a BeH8-mCherry reporter virus, where mCherry expression is decoupled from the viral polyprotein *via* a 2A peptide (39) and flanked by tandem capsid segments (C1 and C2). This design preserves viral replication while enabling real-time tracking of translation efficiency (see *Supplementary Material Figure S2* for construct details).

### Plasmid and vectors

GFP-sensor constructs (GFP-AG GFP-AA) were generated in plenti-PGK with Puro^R^ Vigene Biosciences, Charles River, USA). Corresponding VSV-G pseudo-typed virus-like-particles were produced in HEK293T cells as previously described (40). Additional constructs of GFP sensors were generated to investigate GFP sensor translation after transfection of their corresponding mRNAs. In addition to GFP-AG and GFP-AA, GFP-AAR was synthesized with all six Arg codons (CGC) mutated in AGA U_34_-sensitive codon. Codons and amino acids sequence comparisons of the three GFP variants are given in *Supplementary Material Figure S1 and S4*. All three GFP sensors were subcloned *via* PCR in pGEM-9Zf(-) with a consensus Kozak sequence upstream the Start codon (GCCACC<u>ATG</u>). By using a neural network prediction tool (41, 42) of splice sites in human genes, NetGene2 (https://services.healthtech.dtu.dk/services/NetGene2-2.42), the analysis of the individual GFP cds (-AG, -AA and AAR) and GFP-AG and GFP-AA in plenti-PGK plasmid revealed no cryptic donor or acceptor splice sites within the sequences (*Supplementary Material Figure S13*). Each GFP sensor mRNA (GFP-AG, GFP-AA, or GFP-AAR) was co-transfected with an equal amount of BFP (blue fluorescent protein) mRNA—generated *via* PCR cloning in pGEM-9Zf(-) with a Kozak sequence and derived from plasmid pKLV-U6gRNA(BbsI)-PGKpuro2ABFP (Addgene #50946)—to serve as an internal control for normalizing GFP fluorescence and correcting for variability in transfection efficiency and mRNA expression levels.

### Reagents and antibodies

RECTAS was produced as previously described (43, 44) and diluted in DMSO to treat cells at the required concentration. Anti-ELP1 rabbit polyclonal antibody (#A10127) was from ABclonal. Anti-ALKBH8 rabbit polyclonal antibody (#OACA06811) was from AvivaSysBio. Anti-CTU1 rabbit polyclonal antibody (#9663252) was from Affinity Biosciences. HRP-conjugated Goat-anti-Rabbit antiserum was from Pierce (Thermo Fischer). HRP-conjugated anti-actin was from Sigma-Aldrich.

### *In vitro* transcription of GFP sensors

GFP and BFP cDNA sequences previously subcloned into pGEM-9Zf(-) were submitted to *in vitro* SP6 transcription using mMESSAGE mMACHINE™ SP6 transcription kit (Thermo Fischer Scientific) after vector linearization with adequate restriction enzyme cleavage downstream GFP stop codon. To enhance stability of transfected mRNAs, 3’-end polyadenylation was performed using the Poly(A) Tailing Kit (Thermo Fischer Scientific). Purified poly(A)+ mRNAs were then transfected to HeLa cells by using Lipofectamine™ MessengerMAX™ Transfection Reagent (Thermo Fischer Scientific).

### Codon analysis

Codon adaptation index (CAI) and Relative Synonymous Codon Usage (RSCU) calculations were done using CAIcal tools (http://genomes.urv.es/CAIcal) (45) as previously described (46). Codon frequencies were calculated with Codon Utilization Tool (CUT) website (http://pare.sunycnse.com/cut/index.jsp) (47). Relative synonymous codon usage (RSCU) is defined as the ratio of the observed frequency of codons to the expected frequency given that all the synonymous codons for the same amino acids are used equally. The codon adaptation index (CAI) was used to estimate the extent of bias towards codons that were known to be preferred in highly expressed genes for the given host organism. Cluster analyses were done using Genesis 1.8.1(48) and Cluster 3.0 (49) and visualization were made with Java Treeview (50). CAIs of selected human genes was calculated as above. Corresponding protein abundances were derived from PaxDb database version 4.1 (https://pax-db.org/) (51). Values are given in Supplementary Table 2. Phylogenic tree was generated using EMBL-EBI online “Simple phylogeny” tool (52) (https://ebi.ac.uk/jdispatcher/phylogeny/) with the virus sequences displayed in Supplementary Material Table 1, previously aligned with T-Coffee algorithm (53) (https://ebi.ac.uk/jdispatcher/msa/tcoffee). Analysis of U_34_ codon stretches were performed on ZIKV PF13 sequence. Briefly, all 3423 codons were ordered in a csv file. Consecutive U_34_ codons (K^*AAA*^, Q^*CAA*^, E^*GAA*^ and R^*AGA*^) were recovered using *dplyr* package and plotted with *ggplot2* in R environment. Tables presented in *Supplementary Material Table 3* were generated using *dplyr* tools.

### Flow cytometry and fluorescence microscopy

For the VLP transduction experiments (Figure 3A and *Supplementary Figure S3*), all conditions were performed in triplicate. Flow cytometry was executed using a NovoCyte ACEA flow cytometer (Agilent, MRI platform, CNRS Montpellier). Cells were seeded in 96-well microplates at a density of 20,000 cells per well, and 10,000 events were acquired per sample for each biological replicate. Data were processed using NovoExpress software (Agilent), and the resulting Mean Fluorescence Intensity (MFI) values were used for all subsequent quantitative analyses.

For experiments based on the mRNA transfection model (Figure 3B and 7E), all conditions were performed in triplicate. Flow cytometry was performed on LSR FORTESSA flow cytometer (Becton Dickinson, MRI platform, CNRS Montpellier). Cells were seeded in 24-well microplates at a density of 75,000 cells per well, and 20,000 events per sample were recorded for each biological replicate. Data were analyzed using either the MFI global method (as above) or a custom single-cell analysis pipeline implemented in R (4.5.2) using the *flowCore* package. In the single-cell analysis, fluorescence data (mCherry, GFP, or BFP) were extracted for each individual cell of each fcs sample. GFP signals were normalized to BFP within each cell, and cells were categorized based on their mCherry status (infected or non-infected). The average of BFP-normalized GFP signals was then calculated for both infected and non-infected populations, and these values were used to generate the bar plots presented in Figure 3B and 7E. Both the MFI global method and the single-cell analysis yielded consistent and comparable results.

Fluorescence microscopy images were acquired on a Zeiss AxioImager Z2 microscope (MRI platform) and processed for analysis with ImageJ software (54).

### Quantification of tRNA Modifications by Mass Spectrometry (LC-MS/MS)

RNA preparations enriched in tRNAs were obtained using mirVana™ miRNA Isolation Kit (Thermo Fischer). Fractionation accuracy was checked by capillary electrophoresis on a LabChip GX II (Caliper Life Sciences, Hopkinton, USA) (Supplementary Figure S6) showing that tRNAs consistently represented >75% of the small RNA fraction (Small RNA) with minimal contamination from larger species (5S, 18S, or 28S rRNA). tRNA-enriched samples were enzymatically hydrolyzed to ribonucleosides using benzonase (Santa Cruz Biotech) and nuclease P1 (Sigma) in 10 mM ammonium acetate pH 6.0 and 1 mM MgCl2 at 40 °C for 1 h. Ammonium bicarbonate was then added to a final concentration of 50 mM together with phosphodiesterase I and alkaline phosphatase (Sigma), and further incubated at 37 °C for 1 h. Samples were precipitated with 3 volumes of acetonitrile and supernatants lyophilized and reconstituted in internal standard (I.S.) solution for LC-MS/MS analysis. An Agilent 1290 Infinity II UHPLC system with an ZORBAX RRHD Eclipse Plus C18 150 x 2.1 mm (1.8 μm) column protected with an ZORBAX RRHD Eclipse Plus C18 5×2.1 mm (1.8 µm) guard (Agilent) was used for chromatographic separation of nucleosides. The mobile phase consisted of A: water and B: methanol (both added 0.1% formic acid) at 0.2 ml/min, starting with 5% B for 0.5 min, 5-15% B over 3.5 min, 15-95% B over 3 min, 95% B for 0.5 min, and finally 4 min re-equilibration at 5% B. Mass spectrometric detection was performed using an Agilent 6495 Triple Quadrupole system monitoring the mass transitions 317.1-185.1 (mcm5U), 333.1-201.1 (mcm5s2U), 333.1-183.1 (R-/S-mchm5U), 302.1-170.1 (ncm5U), 298.1-166.1 (m1G, m2G), 268.1-136.1 (A), 284.1-152.1 (G), 244.1-112.1 (C), 245.1-113.1 (U), 264.1-127.1 (13C5-m5U, I.S.), 301.1/152.1 (d3-Gm, I.S.), 246.1/114.1 (d2-U, I.S.), 247.1/115.1 (d2-C, I.S.), and 273.1/136.1 (13C5-A, I.S.), in positive ionization mode (55). The specificity of the observed modifications to the tRNA pool was further validated by analyzing the corresponding Large RNA fraction for all experimental conditions, which showed a total absence of detectable U_34_ modification peaks (mcm^5^U, mcm^5^s^2^U, *R* or *S* mchm^5^U, and ncm^5^U), thereby ruling out interference from high-molecular-weight species such as ribosomal RNAs. For quantification purposes, levels of tRNA U_34_ modifications were normalized to the cumulative levels of reference modifications found exclusively on tRNAs m^1^G, m^2^G (Figure 4A)). Conversely, the SH-SY5Y data presented in Figure 4B and the human primary astrocytes data shown in Supplementary Figure S11 were derived from an independent dataset in which only m^2^G was quantified; baseline normalization for these panels was adjusted accordingly.

### Gene disruption by CRISPR-Cas9 and shRNA

The plasmids for CRISPR-Cas9 were obtained from the Montpellier Genomic Collection Platform (Biocampus, Montpellier, France). Guide RNA targeting U_34_-related genes were designed using three online gRNA-optimizing softwares: CRISPR design (http://crispr.mit.edu.insb.bib.cnrs.fr), CRISPR RNA Configurator (http://dharmacon.gelifesciences.com/gene-editing/edit-r/custom-crrna), and CRISPR gRNA Design tool (https://www.dna20.com/eCommerce/cas9). Guides used in this study are presented in Table I. Guides were cloned into pUC57 attB U6 gRNA vectors (56). The generated plasmid pUC57 attB U6 gRNA was transfected into HeLa cells with Lipofectamine 2000, along with the pSpCas9(BB)-2A-GFP (PX458) plasmid (57). At 6 h after transfection, the cells were trypsin treated and resuspended in complete DMEM medium at 2 × 10^4^ cells per ml. Portions (200 μl) of the cell suspension (4 × 10^3^ cells) were used to inoculate 96-well plates and to isolate single cell-derived clones by serial dilution. Isolated green fluorescent protein (GFP)-positive clones were amplified and analyzed by Western blotting to check target gene expression. ShRNAs were Mission shRNAs from SIGMA cloned into plKO.1-puro presented in Table II. Corresponding VSV-G pseudo-typed virus-like-particles were produced in HEK293T cells as previously described (40) and used to transduce A549 cells. Individual clones were isolated after puromycin selection and analyzed by Western blotting to check target gene expression.

**Table I.**
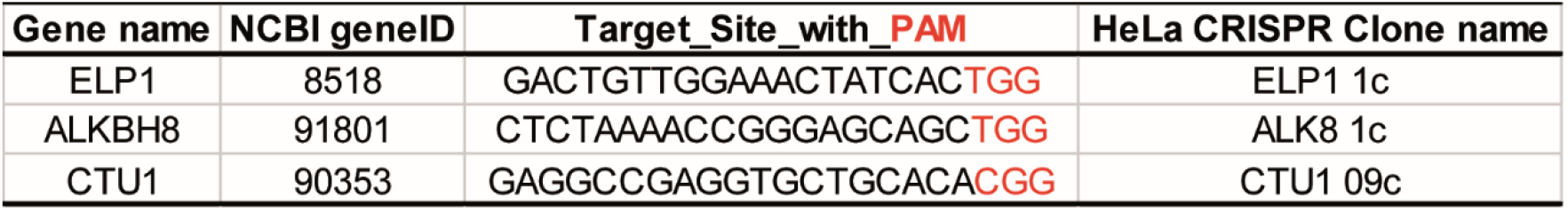
CRISPR/Cas9 guides targeting U_34_ enzyme genes.

**Table II.**
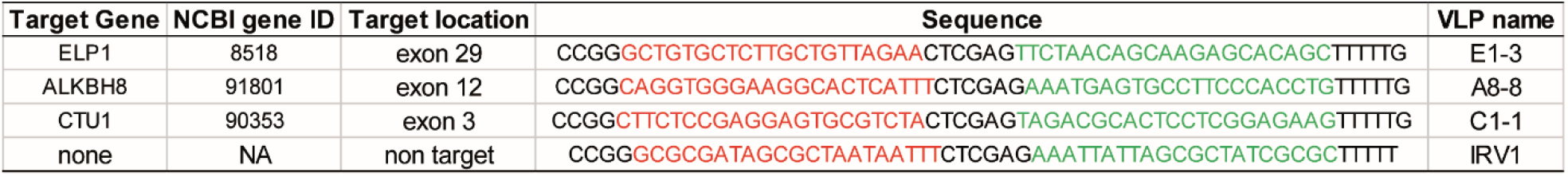
shRNAs targeting U_34_ enzyme genes.

### Assays for Viral Replication

Cells were lysed with the Luna cell-ready lysis module (New England Biolabs). The amplification reaction was run on a LightcyclerR 480 thermocycler (Roche Diagnostics) using the Luna Universal One-Step RT-qPCR kit (New England Biolabs), and ZIKV_For: 5′-AGGATCATAGGTGATGAAGAAAAGT (hybridizes at the end of NS5 sequence); ZIKV_Rev: 5′-CCTGACAACATTAAGATTGGTGC (hybridizes in the 3’UTR region); GAPDH_For: 5′-GCTCACCGGCATGGCCTTTCGCGT and GAPDH_Rev: 5′-TGGAGGAGTGGGTGTCGCTGTTGA primers. Each RTqPCR was performed from biological triplicates, and the means and standard deviations were calculated. Relative quantification of data obtained from RT-qPCR was used to determine changes in ZIKV gene expression across multiple samples after normalization to the internal reference GAPDH gene. GAPDH and ZIKV amplicons are respectively 200 bp and 116 bp in length.

## Results

### ZIKV genome is heavily biased toward A-ending codons

To assess the extent of codon usage bias (CUB) in ZIKV genomes, we performed a comprehensive bioinformatic analysis, including calculations of the Codon Adaptation Index (CAI) and Relative Synonymous Codon Usage (RSCU). These analyses suggest that ZIKV translation efficiency in host cells may be significantly influenced by the availability of compatible tRNAs (Figure 2). Global CAI analysis revealed that none of the ZIKV genomes examined, including the ancestral African strain MR766 and the Asian strains PR15, BeH8, PF13 or SPH15, are fully adapted the human or mosquito (*Aedes aegypti*) host codon preference, with CAIs between 0.755 and 0.759 relative to *Homo sapiens* and between 0.728 and 0.733 relative to *Aedes aegypti* (Figure 2A and Supplementary Material Table 1). For comparison, highly expressed human genes with protein abundances exceeding 10,000 ppm, such as β-globin, β-myosin or β-tubulin, exhibit higher CAI values (0.869, 0.882, and 0.819 respectively) (Supplementary Material Table 2). In contrast, poorly expressed human genes, such as RHA, RIG-I or Kallmann Syndrome protein, with protein abundances below 300 ppm, show lower CAI values (0.748, 0.765 and 0.778, respectively). For further analysis, we selected the Asian strain PF13 as a representative prototype due to its phylogenetic clustering with other Asian strains (SPH15, PR15, BeH8). By mapping CAI scores across the PF13 genome using *Homo sapiens* codon usage as a reference, we observed a non-uniform distribution of translational adaptation (Figure 2B). Several stretches in the ZIKV genome are characterized by poorly adapted codons, resulting in CAI values between 0.5 and 0.6, potentially representing translational bottlenecks in the host.

**Figure 2.**
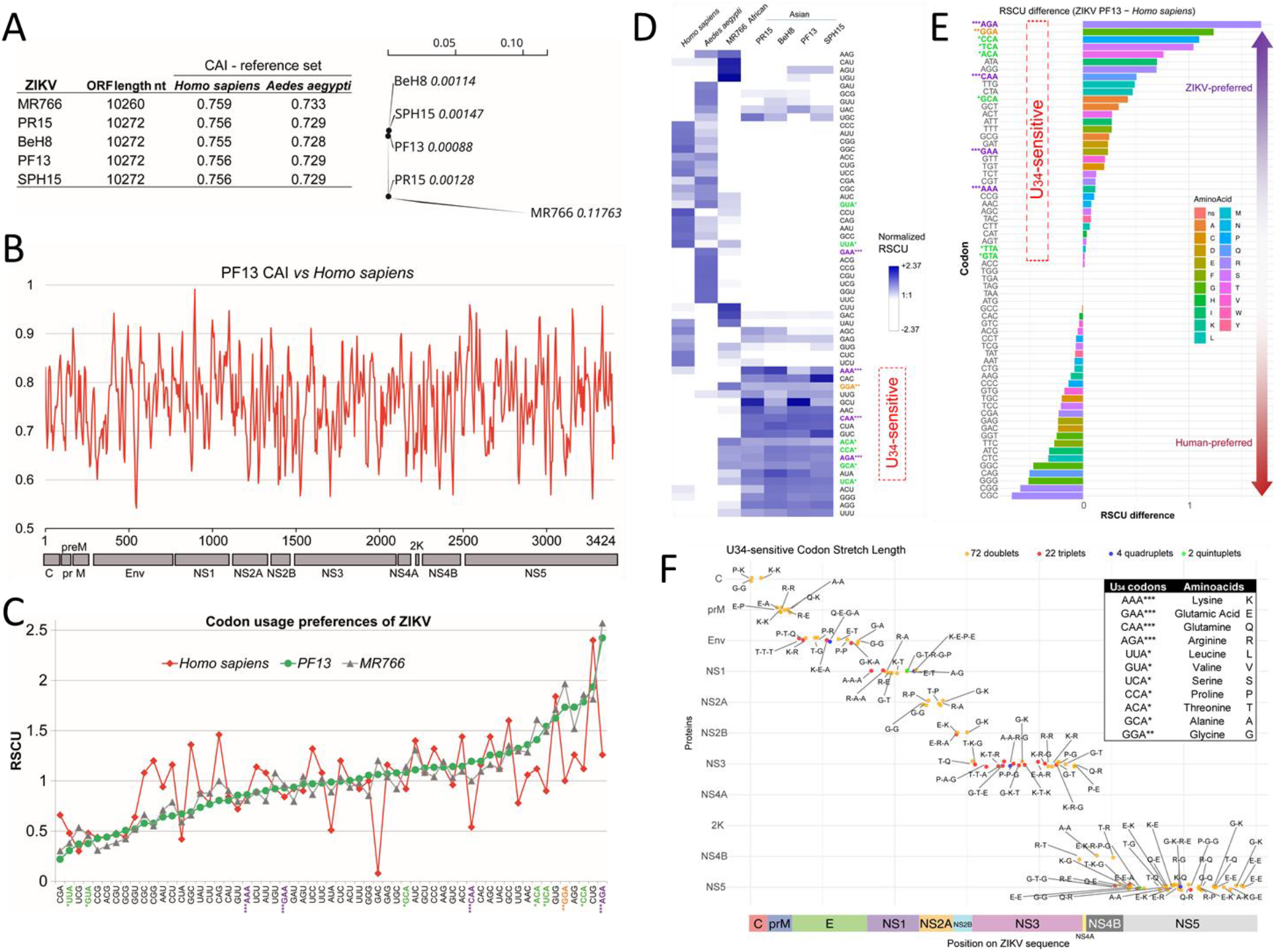
Codon usage and preferences of ZIKV. (A) Global CAI values for various ZIKV strains compared to *Homo sapiens* reference codon usage alongside a phylogenetic analysis showing the divergence between African and Asian ZIKV strains (scale: genetic distance). (B) CAI profile plotted along of the genome of the ZIKV PF13 strain (Asian genotype isolate from French Polynesia 2013) relative to *H. sapiens* reference codon usage preference. (C) Comparative analysis of Relative Synonymous Codon Usage (RSCU) values for ZIKV strains MR766 and PF13, and for *H. sapiens*. Codons are ordered according to their RSCU values for the PF13 strain. (D) Hierarchical clustering of normalized RSCU values across different ZIKV genotypes compared to *Homo sapiens* and *Aedes aegypti* coding sequences. Agglomeration was performed using the average linkage method (WPGMA: weighted pair group method with averaging). (E) RSCU differences between ZIKV PF13 and *Homo sapiens*, ranked from human-preferred (bottom) to ZIKV-preferred codons (top). U_34_-sensitive codons are highlighted with their corresponding U_34_ modifications (*: ncm^5^U in green, **: mcm^5^U in orange and ***: mcm^5^s^2^U in purple). (F) Mapping and frequency of U_34_-sensitive codon stretches across the ZIKV genome. Each point represents a cluster of consecutive U_34_-sensitive codons (K^*AAA*^, Q^*CAA*^, E^*GAA*^ and R^*AGA*^), plotted according to its position on the ZIKV sequence (*x-axis*) and the specific viral protein in which it occurs (*y-axis*). The color of each point indicates the length of the codon repeat and labelled with the specific amino acid sequence (motif) resulting from the codon stretch. The inset table lists the specific U_34_-sensitive codons analyzed and their corresponding amino acids. Refer to *Supplementary material Table 3* for a complete list of codon positions, specific sequences, and gene assignments.

RSCU analysis further confirmed a mismatch between ZIKV PF13 codon usage and the codon preference of *Homo sapiens* (Figure 2C). Codons highly enriched in ZIKV are infrequently utilized in the human genome. Suboptimal codons, defined as codons that are less efficiently decoded due to limited availability of cognate tRNAs or reliance on specific tRNA modifications, were particularly prominent in this analysis. Hierarchical clustering based on RSCU values revealed a stark divergence in codon preference between the viral strains and their hosts (Figure 2D). Both the African (MR766) and Asian (PR15, BeH8, PF13, SPH15) ZIKV lineages showed a concentrated enrichment for a specific cluster of A-ending codons while human and *Aedes aegypti* genomes favor G-ending codons. Notably, the viral-preferred codons were heavily enriched for U_34_-sensitive codons, which require specific chemical modifications at the tRNA wobble position for efficient translation. Specifically, codons such as AAA, CAA, and AGA (marked with ***) showed high normalized RSCU values across all viral strains, particularly within the Asian lineage. This viral preference suggests that ZIKV may be particularly sensitive to the host’s tRNA modification landscape. Furthermore, the Asian lineage strains displayed a highly conserved and uniform RSCU profile, suggesting that the recent evolution of ZIKV has maintained a stable codon bias that distinguishes it from its hosts and, to a lesser extent, the ancestral African lineage. While the ancestral African genotype (MR766) is distinct from both the ZIKV Asian genotypes and the human genome, it also shows a light enrichment in U_34_-sensitive codons (Figure 2D). Of note, we noticed an enrichment of Glu^*GAA*^ in *Aedes aegypti* suggesting that this codon is likely optimized for the mosquito’s tRNA pool, which may include cognate tRNAs with mcm^5^s^2^U_34_ modifications that promote efficient decoding of Glu^*GAA*^. In contrast, *Homo sapiens* coding sequences do not show this enrichment, indicating a potential mismatch in tRNA availability for this codon between the two hosts.

To characterize the translational landscape of ZIKV, we calculated the RSCU difference between the ZIKV strain PF13 and the *Homo sapiens* genome (Figure 2E). This analysis confirmed a significant divergence in codon usage preferences, where codons highly enriched in ZIKV are infrequently utilized in the human genome. Notably, the viral-preferred codons were heavily enriched for U_34_-sensitive codons. Specifically, codons such as AGA (***), CAA (***), GAA (***), and AAA (***) showed the highest positive RSCU differences. Other U_34-_sensitive codons, including GGA (**), CCA (*), and ACA (*), were also significantly enriched in the virus relative to the host.

The clustering of suboptimal codons into “stretches” (consecutive sequences) has been proposed as a mechanism that increases a gene’s sensitivity to changes in the host epitranscriptome and tRNA modification levels (14, 58). When U_34_ tRNA modifications are impaired, suboptimal codon pairs of U_34_-sensitive codons are sufficient to trigger ribosome collisions and alter local translation dynamics (59). Our analysis of the spatial arrangement of U_34_-sensitive codons across the entire ZIKV polyprotein revealed a high frequency of clustered U_34_-sensitive codons throughout the ZIKV genome, identifying 72 doublets, 22 triplets, 4 quadruplets, and 2 quintuplets (Figure 2F). These stretches are distributed across all ZIKV proteins, including the structural proteins (C, prM, and Env) and the non-structural proteins (NS1 through NS5). While U_34_-sensitive codon pairs are distributed widely throughout the polyprotein, higher-order clusters—such as triplets, quadruplets and quintuplets—are specifically concentrated within essential non-structural enzymes like NS1, NS3 and NS5. These high-density clusters are located in enzymes critical for the early stages of the viral life cycle, highlighting specific genomic regions that may act as translational bottlenecks.

Collectively, these findings suggest that optimal translation of ZIKV proteins in human cells may require a profound reprogramming of the host tRNA pool, particularly targeting tRNAs that require U_34_ modification, to accommodate the virus’s highly skewed codon usage.

### ZIKV infection enhances translation of A-ending codons

To experimentally assess whether ZIKV infection enhances the translation of A-ending codons, we took advantage of two engineered eGFP reporter constructs with opposing codon bias for the three amino acids that require mcm^5^s^2^U_34_ modification on their respective tRNAs for efficient decoding of AA-ending codons. The AG-biased eGFP (GFP-AG) corresponds to the human-optimized eGFP sequence in which 42 out of 44 codons for Lys, Gln or Glu, are AG-ending, reflecting the codon usage of highly translated human genes. Conversely, the AA-biased GFP (GFP-AA) construct is a fully synthetic variant in which all such codons have been changed to their AA-ending counterparts (Figure 3A, left), thereby mimicking the codon preference observed in the ZIKV genome (a sequence alignment of both GFP constructs is provided in *Supplementary Material Figure S1*). Both GFP sensors were delivered by lentiviral vectors in ZIKV-permissive HeLa cells pre-infected with BeH8-mCherry, a modified ZIKV strain engineered to express the mCherry marker (*Supplementary Material Figure S2)*, as a proxy of ZIKV translation efficiency allowing identification of infected cells. GFP expression was preliminary monitored at different time points after transduction (*Supplementary Material Figure S3*) using both fluorescence microscopy (*Figure S3B*) and flow cytometry (*Figure S3C*) to evaluate their relative translation efficiency under ZIKV infection. Immunofluorescence analysis, conducted 72 hours post-transduction with codon-biased GFP sensors, revealed a dramatic shift in translational efficiency upon ZIKV infection. Specifically, there was a marked increase in GFP-AA expression alongside a concurrent reduction in GFP-AG expression. Quantification of mean fluorescence intensity per square pixel confirmed that, in non-infected cells, GFP-AG was translated more efficiently than GFP-AA, consistent with the codon preferences of human cells. In contrast, ZIKV-infected cells exhibited the opposite pattern, with GFP-AA expression surpassing that of GFP-AG (Figure S8C, right panel). This shift was further validated by flow cytometry analysis of ZIKV-infected mCherry+ cells (Figures S8D and S8E S3D and S3E).

**Figure 3.**
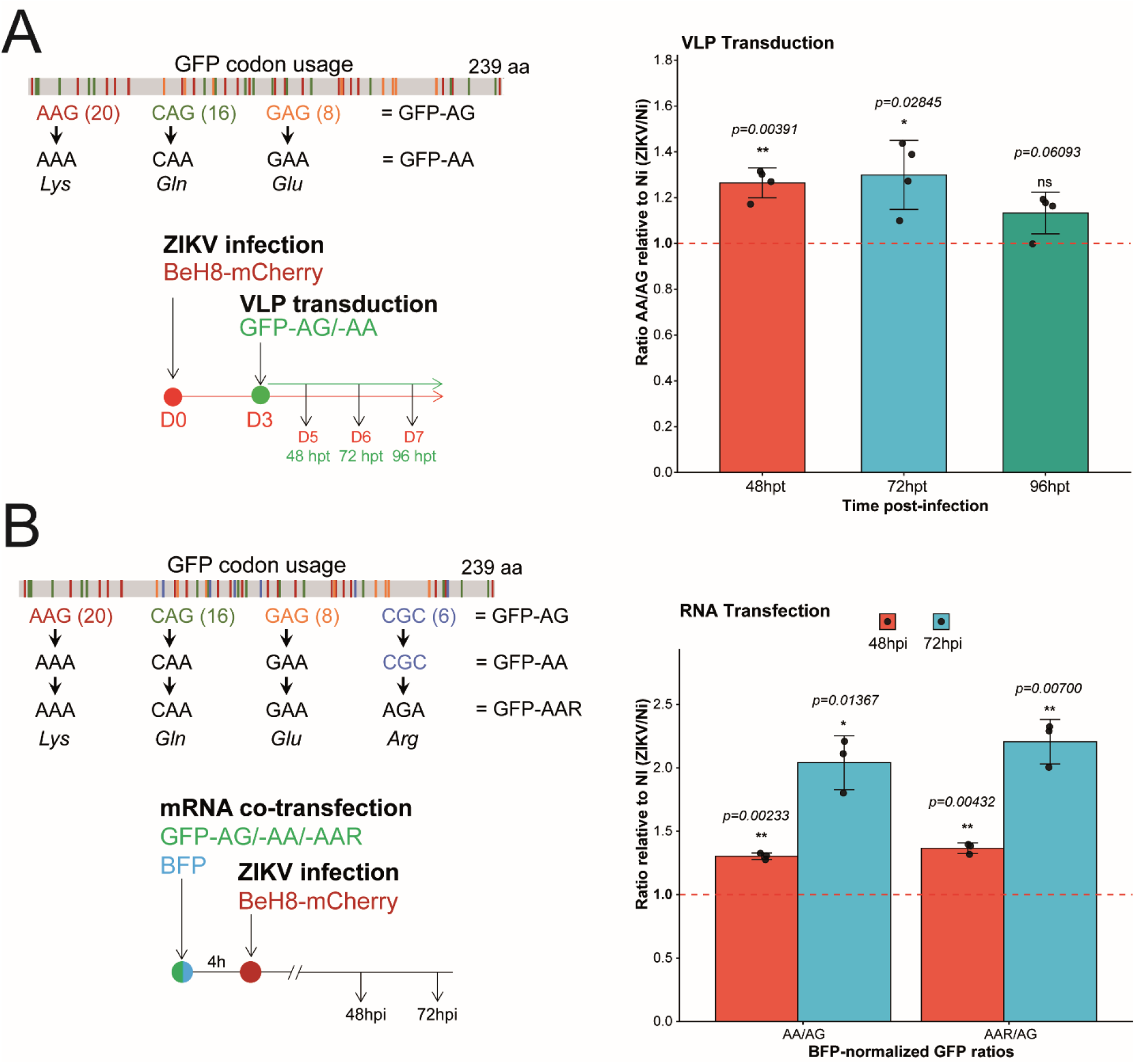
Impact of ZIKV infection on codon-specific translation. A. VLP Transduction Model. Schematic of GFP-AG and GFP-AA constructs, differing in synonymous codon usage for Lys, Gln, and Glu and experimental workflow (left). Ratio of GFP-AA to GFP-AG expression in ZIKV-infected cells relative to non-infected cells at 48, 72, and 96 hours post-transduction (hpt). B. RNA Transfection Model. Schematic of GFP-AG, GFP-AA, and GFP-AAR constructs. GFP-AAR includes six U_34_-sensitive Arg codons in addition to A-ending codons for Lys, Glu, and Gln, and strategy of transfection/infection in this model (left). BFP-normalized GFP ratios (AA/AG and AAR/AG) in ZIKV-infected cells relative to non-infected cells at 48- and 72-hours post-infection (hpi) (right).

To better visualize this phenomenon, we used the GFP-AA/GFP-AG ratio in ZIKV-infected cells relative to uninfected cells. This ratio serves as a proxy for how ZIKV infection enhances the expression of its codon-biased genome (mimicked by GFP-AA) at the expense of host genes (reported by GFP-AG) (Figure 3A, right). At early time points post-transduction, we observed a transient increase in the GFP-AA/GFP-AG expression ratio, indicating a preferential translation of AA-ending codons. This effect progressively attenuated over time, with the AA/AG ratio returning near baseline levels.

To complement the previous VLP transduction model, we developed an alternative approach involving transfection of GFP sensor mRNAs followed by infection with ZIKV BeH8-mCherry. Given the high prevalence of Arg^*AGA*^ codons sensitive to U_34_ modification in the ZIKV genome (Figure 2C and 2E), we also generated an additional GFP sensor variant, GFP-AAR (*Supplementary Material Figure S4*), which incorporates six U_34_-sensitive Arg synonymous codons alongside the A-ending Lys, Glu, and Gln codons already present in GFP-AA (Figure 3B, left). To control for cell-to-cell variability in transfection efficiency, we co-transfected cells with a BFP mRNA alongside each GFP mRNA. This enables precise normalization of GFP fluorescence intensities, ensuring robust quantification of the translational changes induced by ZIKV infection. By comparing the GFP-AA/GFP-AG and GFP-AAR/GFP-AG ratios in ZIKV-infected cells relative to non-infected controls (Figure 3B, right), we observed a significant and sustained increase in the translation of GFP sensors containing AA-ending or AA-ending plus rare Arg^*AGA*^ codons during infection. Notably, the GFP-AAR construct shows at 72hpi an AAR/AG ratio slightly higher than the AA/AG, suggesting a high contribution of Arg^*AGA*^ codons in addition to AA-ending codons (Lys^*AAA*^, Gln^*CAA*^ and Glu^*GAA*^) to the ZIKV codon-dependent translation. This approach not only controlled for variability in transfection efficiency but also provided a clear and quantitative measure of how ZIKV infection reprograms host translation machinery to favor viral protein synthesis.

These findings suggest that ZIKV infection transiently alters host translation machinery, favoring the decoding of A-ending codons (Lys, Gln, Glu, and Arg) while suppressing AG-ending codon translation. This translational shift, evidenced by elevated GFP-AA and/or GFP-AAR expression relative to GFP-AG, is consistently observed in both VLP transduction and RNA transfection models, highlighting a potential viral strategy to optimize its own protein synthesis at the expense of host gene expression.

### Dynamic variation of tRNA modifications in ZIKV-infected cells

Given the enhanced translation of the GFP-AA and GFP-AAR sensors observed during ZIKV infection, likely dependent on the host cell’s ability to decode A-ending codons through specific tRNA modifications, we investigated the dynamic changes in tRNA post-transcriptional modifications (PTMs) in ZIKV-infected brain cells. These cells, known to be highly permissive to ZIKV, offer a physiologically relevant model for studying neurotropic virus-host interactions. We infected two types of human brain-resident cells: the neuroblastoma SH-SY5Y cell line, and human primary astrocytes. Following infection, we confirmed viral replication by RT-qPCR (Supplementary Figure S5). Total RNA was extracted and fractionated to isolate small RNA species, in which tRNA represent over 75 % (Supplementary Figure S2 S6).

We then investigated how ZIKV infection altered the abundance of individual PTMs. LC-MS/MS analysis of SH-SY5Y and astrocytic cells identified more than 20 distinct tRNA modifications, several of which were significantly modulated by ZIKV infection (data not shown). We then focused specifically on the critical U_34_-based modifications, revealing that mcm^5^U and mcm^5^s^2^U were clearly increased in SH-SY5Y cells at 72 hours post infection (Figure 4A). Notably, mcm^5^s^2^U levels were enhanced in the early stages of infection (Figure 4B), peaking between 24 and 72 hpi, followed by a progressive decline at 96 hpi, although viral production was increasing. This pattern mirrors the temporal dynamics of GFP-AA translation enhancement observed during ZIKV infection (Figure 3A), suggesting that dynamic regulation of mcm^5^S^2^U may contribute to the translational reprogramming induced by the virus. We performed the same analysis on primary human astrocytes and observed a remarkably similar trend, characterized by a transient, early increase in mcm^5^s^2^U levels at 48 hpi followed by a decline at 120 hpi (*Supplementary Figure S9*). However, due to the high sensitivity of primary astrocytic cells to ZIKV infection, their use in this context introduced certain reproducibility challenges that warrant further verification.

**Figure 4.**
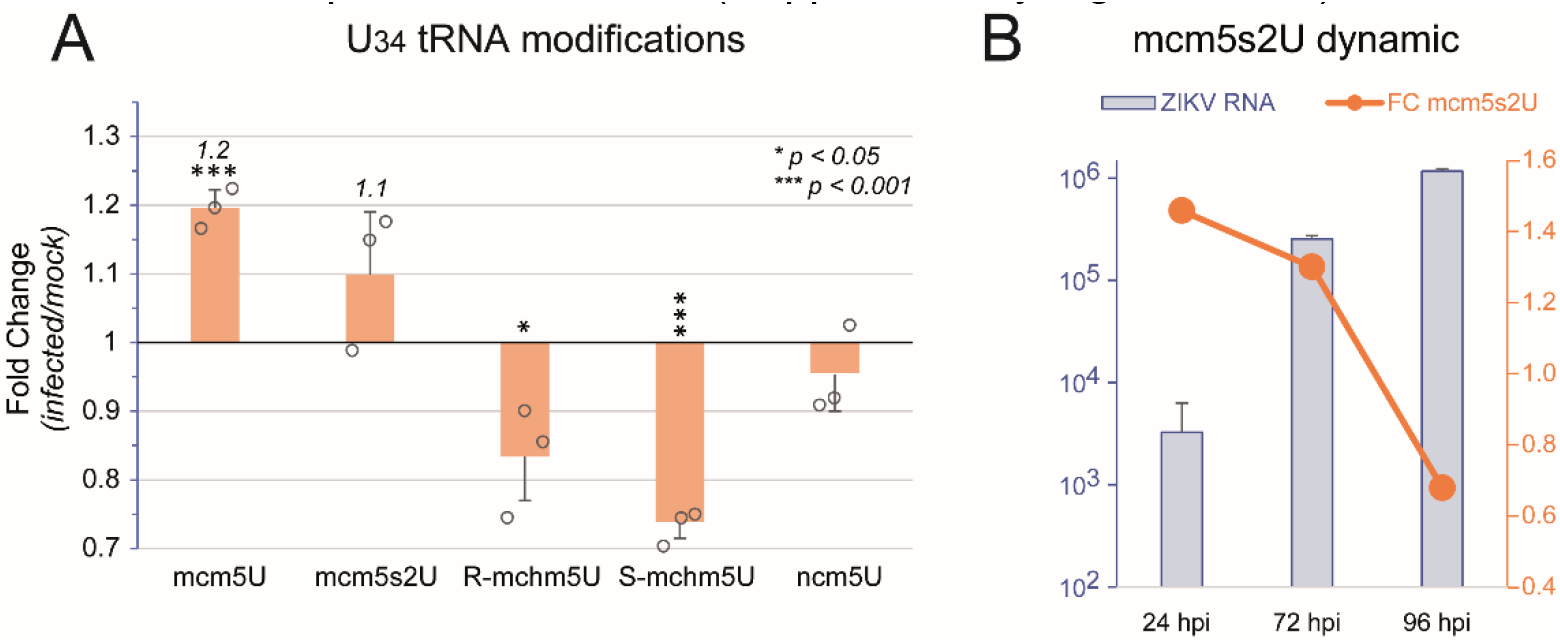
Modulation of tRNA U_34_ modifications during ZIKV infection. (A) Fold change of specific tRNA U_34_ modifications in infected SH-SY5Y cells 72 hours post infection relative to mock controls. Data represent the mean fold change +/-standard error of the mean (SEM) for the indicated chemical. Individual data points are plotted as open circles. Data were normalized to the cumulative levels of reference modifications found exclusively on tRNAs m1G and m2G. Corresponding original chromatograms are presented in *Supplementary Figure S7*. (B) Temporal dynamics of mcm5s2U levels in relation to ZIKV RNA replication. ZIKV RNA levels and the corresponding fold change (FC) of the mcm^5^s^2^U modification were tracked at 24, 72, and 96 hours post-infection (hpi). Error bars indicate SEM. Corresponding original chromatograms are presented in *Supplementary Figure S8*.

While these global shifts of global fold change of mcm^5^U and mcm^5^s^2^U would appear modest (ranging from 1.1 to 1.4), it masks a substantial, targeted increase in the mcm^5^U34/mcm^5^s^2^U34-modifiable tRNA subset (GlnUUG, LysUUU, GluUUC, ArgUCU), which constitutes just 5.6–8.2% of the total tRNA pool. This proportion is corroborated by human tRNA gene diversity analyses (60) and validated across mim-tRNAseq (61), MarathonRT-tRNAseq (62), Induro-tRNAseq (63), and nanopore-based tRNAseq (64, 65). When adjusted for this subset, the localized fold-change reaches 3.4-to 4.5-fold, revealing a non-linear, inverse relationship between the fraction of modifiable tRNAs in the pool and their specific fold-change. As illustrated in Supplementary *Fig S10*, smaller fractions of modifiable tRNAs yield exponentially higher localized effects, demonstrating that even minor global increases in mcm^5^U/mcm^5^s^2^U modifications can signify a major impact on this critical subset. This selective amplification underscores the targeted enhancement of mcm^5^U/mcm^5^s^2^U modification in tRNAs decoding wobble U_34_-dependent codons, suggesting a mechanism by which ZIKV may prioritize translation of transcripts enriched in these codons.

### ZIKV infection efficiency depends on the functional integrity of the U_34_-tRNA modification pathway

In mammals, the enzymes responsible for catalyzing chemical U_34_ modification include the Elongator complex (ELP1–6), the tRNA methyltransferase ALKBH8 and the thiouridylases CTU1/CTU2 (66). If ZIKV translation indeed depends on the presence of the mcm^5^s^2^U modification at U_34_, then efficient viral replication should require a fully functional Elongator complex. To test this hypothesis, we used primary fibroblasts derived from patients with Familial Dysautonomia (FD), a genetic disorder characterized by loss-of-function mutations in *ELP1* (also known as IKBKAP) (67), resulting in reduced levels of mcm^5^s^2^U tRNA (68). In FD fibroblasts, we observed markedly reduced levels of ZIKV translation and infection compared to control fibroblasts (WT) derived from healthy individuals, as indicated by the diminished expression of the virus-encoded mCherry reporter (Figure 5A). Although fibroblasts are poorly infected by ZIKV (typically 1-5 % infected cells), quantification of mCherry fluorescence intensity serves as a useful proxy for viral translation efficiency. Strikingly, the proportion of mCherry-expressing cells was drastically reduced in ELP1-deficient FD fibroblasts relative to controls, strongly suggesting that Elongator-mediated U_34_ modifications are critical for efficient ZIKV translation (Figure 5B). This phenotype was also observed, though to a lesser extent, across a range of lower MOIs (0.1, 0.5 and 1.0) (Figure 5C), where infection levels were monitored by measuring ZIKV RNA copy number using RT-qPCR, normalized against GAPDH. By monitoring mCherry fluorescence intensities 48 h and 72 h after ZIKV infection at high MOI (20), we observed a strong and consistent reduction in reporter signals in FD cells compared to WT cells (Figure 5D). Given that mCherry fluorescence intensities correlate with ZIKV translation levels (*Supplementary Material Figure S2*), these results strongly support a model in which the absence of functional Elongator-mediated U_34_ modifications disrupts viral protein synthesis. Two independent experiments conducted at MOIs of 10 and 20 with recording fluorescence intensities at 48 h and 72 h post-infection yielded identical results (Supplementary Figure S11), further reaffirming the crucial role of the Elongator complex and Elongator-mediated U_34_ tRNA modifications in facilitating efficient ZIKV infection *in vitro*.

**Figure 5.**
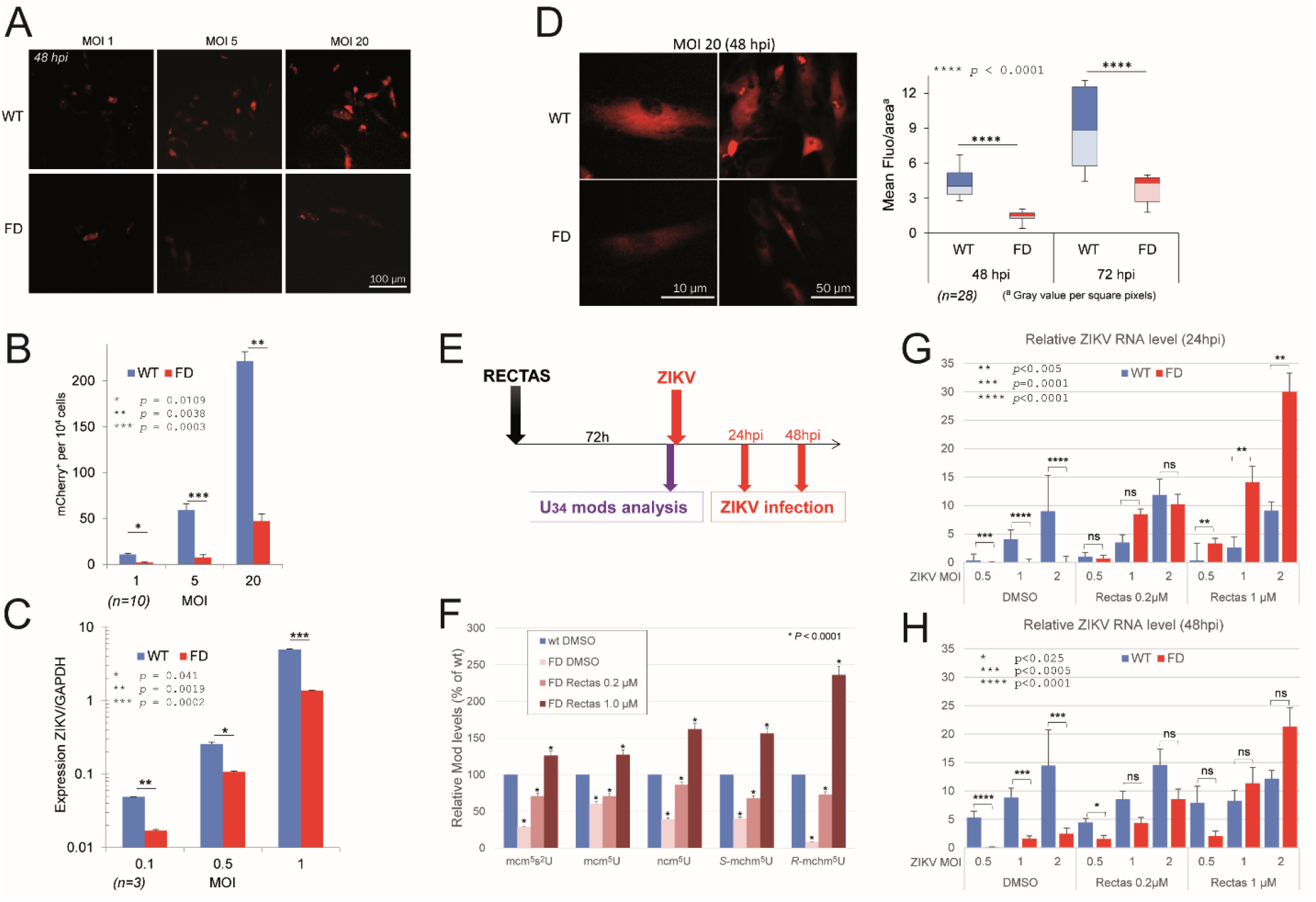
Impaired ZIKV infection in FD fibroblasts lacking U_34_ tRNA modifications can be reverted by RECTAS treatment. (A). Human primary fibroblasts derived from skin biopsies from either a healthy control (WT) or from a patient with Familial Dysautonomia (FD) were infected with ZIKV BeH8-mCherry at MOI of 1, 5 or 20. (B) 48 h post-infection, ZIKV replication was assessed by quantifying the number of mCherry-positive cells per 10^4^ cells using fluorescence microscopy. (C) ZIKV infection levels at low MOIs (0.1. 0.5 and 1.0) were quantified by RTqPCR 48 h post-infection. (D) mCherry fluorescence intensities were visualized by fluorescence microscopy. Fluorescence intensities per square pixel were quantified with ImageJ in non-FD (WT) and FD fibroblasts at 48 h and 72 h post-infection (hpi) at MOI of 20. Results are presented as box-plots where *n* corresponds to the number of cells analyzed. (E) Experimental workflow outlining the use of RECTAS, a small molecule previously shown to upregulate U34 modification pathways, for functional rescue experiments in FD fibroblasts. (F) U_34_ tRNA modification levels in FD cells following 72 h RECTAS treatment (0.2 µM and 1.0 µM) compared to untreated FD cells and WT controls. (G and H), ZIKV infection efficiency in RECTAS-treated WT and FD fibroblasts infected at MOIs of 0.5, 1.0 and 2.0. Viral RNA levels were determined by RTqPCR at 24 hpi (G) and 48 hpi (H) and normalized to GAPDH.

In parallel, we took advantage of the small molecule RECTAS, which is known to rescue ELP1 expression and normal mcm^5^s^2^U_34_ levels(43) in cells from FD patients, to generate a cellular system with graded levels of U_34_-modified tRNAs. FD patients carry a common IVS20 + 6 T > C mutation in the *ELP1* gene that impairs proper recognition of the 5′ splice site by U1 snRNP, resulting in exon 20 skipping, a frameshift, and a premature termination codon in exon 21. This defective splicing event markedly reduces ELP1 expression and impairs tRNA modification (69). RECTAS corrects this splicing defect, restoring normal ELP1 mRNA processing and expression. Functional U_34_ tRNA modifications are subsequently re-established, offering a potential therapeutic avenue for FD patients (44). To determine whether this correction could also restore ZIKV susceptibility, FD and WT fibroblasts were treated with 0.2 µM or 1.0 µM RECTAS for 72 h before ZIKV infection (Figure 5E). Mass spectrometry analysis confirmed that RECTAS treatment efficiently restored five U_34_ tRNA modifications in FD cells to levels comparable to those of WT cells (Figure 5F). Three days after treatment, cells were infected with ZIKV, and infection levels were monitored by RTqPCR-quantification of viral RNA at 24 h and 48 h post-infection. Notably, RECTAS treatment progressively restored ZIKV infection levels in FD fibroblasts, reaching levels similar to WT cells (Figure 5G and 5H). These results demonstrate that pharmacological restoration of U_34_ tRNA modifications reactivates ZIKV protein synthesis in FD cells, reinforcing the critical role of the Elongator-dependent tRNA modification pathway in facilitating efficient viral translation and replication.

### Inactivation of U_34_ tRNA modification enzymes via CRISPR/Cas9 and shRNA impairs ZIKV infection

To further validate the critical role of U_34_ tRNA modifications in ZIKV replication, we disrupted the activity of enzymes responsible for the stepwise biosynthesis of mcm^5^s^2^U-modified tRNAs. Using CRISPR/Cas9, we generated HeLa cells with individual knockouts of *ELP1, ALKBH8* and *CTU1*, each encoding a key enzyme in the stepwise formation of the mcm^5^s^2^U modification (Figure 1B). After clonal selection, KO cells were challenged with ZIKV-mCherry (BeH8-strain) at a MOI of 0.5. The level of infection was then monitored by quantifying mCherry-positive cells between 4- and 26-days post-infection (dpi) (Figure 6A). At 4 dpi, the infection level in ELP1-KO cells was approximately 25 % lower than in the WT control. Over time, viral infection progressive cleared from the U_34_-KO cell population. By 26 dpi, only a few KO cells remained infected (Figure 6B). In contrast, control cells reached 100 % infection at 4 dpi, and succumbed to virus-induced cytotoxic effects between 8 and 15 dpi. Knockout efficiency was validated by Western blot analysis (Figure 6C). These data demonstrate that cells devoid of U_34_ tRNA-modifying enzymes are unable to support the full replication and sustained spread of ZIKV.

**Figure 6.**
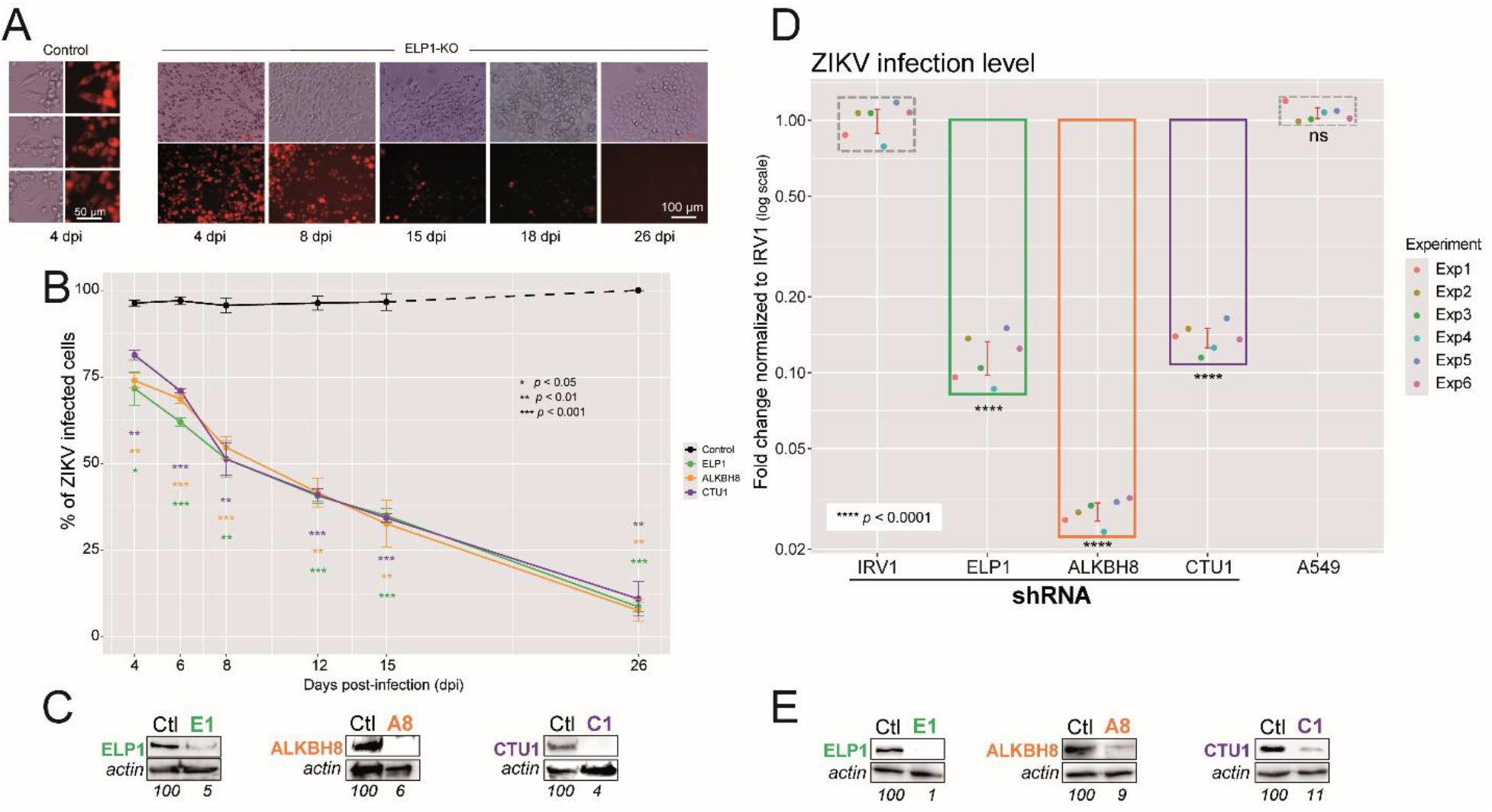
Disruption of the tRNA U_34_ modification pathway impairs ZIKV infection. (A) Representative time course images of ZIKV-infection (MOI = 0.5) in control HeLa cells (left) and ELP1-CRISPR/Cas9 KO cells. Infection was assessed by fluorescence microscopy based on mCherry reporter expression. (B) Quantification of ZIKV infection in controls and CRISPR/Cas9-edited HeLa clones lacking ELP1, ALKBH8 or CTU1. Measurements were conducted in triplicate. (C) Western blot validation of gene knockouts in CRISPR/Cas9-edited HeLa clones. (D) Puromycin-selected A549 cells stably expressing the indicated shRNAs were infected with ZIKV at MOI of 0.5. At 48 h post-infection, viral RNA levels were determined by RTqPCR using GAPDH as an internal control. A549 corresponds to ZIKV-infected cells without shRNAs. Data represent 6 independent experiments (n=6). (E) Western blot analysis of shRNA-treated A549 clones showing protein expression for the respective targeted genes.

In parallel to CRISPR/Cas9, we also employed shRNAs-mediated knockdown to reduce the expression of ELP1, ALKBH8 and CTU1 in human A549 cells. Following lentiviral transduction of shRNAs targeting each gene or a non-targeting irrelevant control (IRV1), and puromycin selection, cells were infected with ZIKV (MOI=0.5), and viral infection was quantified by RT-qPCR quantification. Knockdown of each enzyme in the three steps of U_34_ tRNA modifications (Figure 6E) resulted in substantial reduction of genomic RNA levels, confirming impaired replication (Figure 6D). Together, the results from both gene inactivation consistently corroborate the essential role of mcm^5^s^2^ writers in ZIKV replication. While full knockouts resulted in the progressive clearance of ZIKV infection, partial shRNA knockdowns led to a noticeable attenuation of the infection. Collectively, these complementary experiments provide strong evidence that mcm^5^s^2^ modification of tRNAs is crucial for efficient ZIKV RNA translation and replication.

### Impact of U_34_ modification on codon-dependent ZIKV translation

To validate whether U_34_ tRNA modifications are directly required for ZIKV translation, we infected wild-type (WT) and ELP1-knockout HeLa cells with a ZIKV BeH8-mCherry reporter virus to track translation dynamics in real time (Figure 7A, *Supplementary Material Figure S2*) Importantly, to confirm that the observed effects were specific to U_34_-dependent codon decoding and not due to global translation inhibition, we compared GFP and BFP fluorophore expression in wild-type and ELP1-KO cells. The absence of significant differences in mean fluorescence intensity between these cell types (*Supplementary Material Figure 12*) confirmed that global translation capacity remains unchanged in ELP1-KO cells. After validating the ELP1 knockout (Figure 7B), we observed that ELP1-KO cells displayed significantly reduced mCherry expression (Figure 7D) across all tested multiplicities of infection (MOIs: 0.1, 2, and 5) coincidating with lower infection levels. These results indicate that the absence of U_34_ modifications impairs both ZIKV infection and viral protein synthesis. Using our RNA transfection model of codon-biased GFP sensors and BFP RNA for normalization (as in Figure 3B), we found that the translation of codon-biased GFP constructs (GFP-AA and GFP-AAR) relative to GFP-AG was significantly reduces in ELP1-KO cells compared to wild-type cells (Figure 7E). Notably, the AAR/AG ratio was even more diminished than the AA/AG ratio in ELP1-KO cells, emphasizing the preponderant contribution of Arg^*AGA*^ codons in codon-specific translation. This highlights the critical role of U_34_ modifications in enabling efficient decoding of A-ending and U_34_-sensitive codons during ZIKV infection.

**Figure 7.**
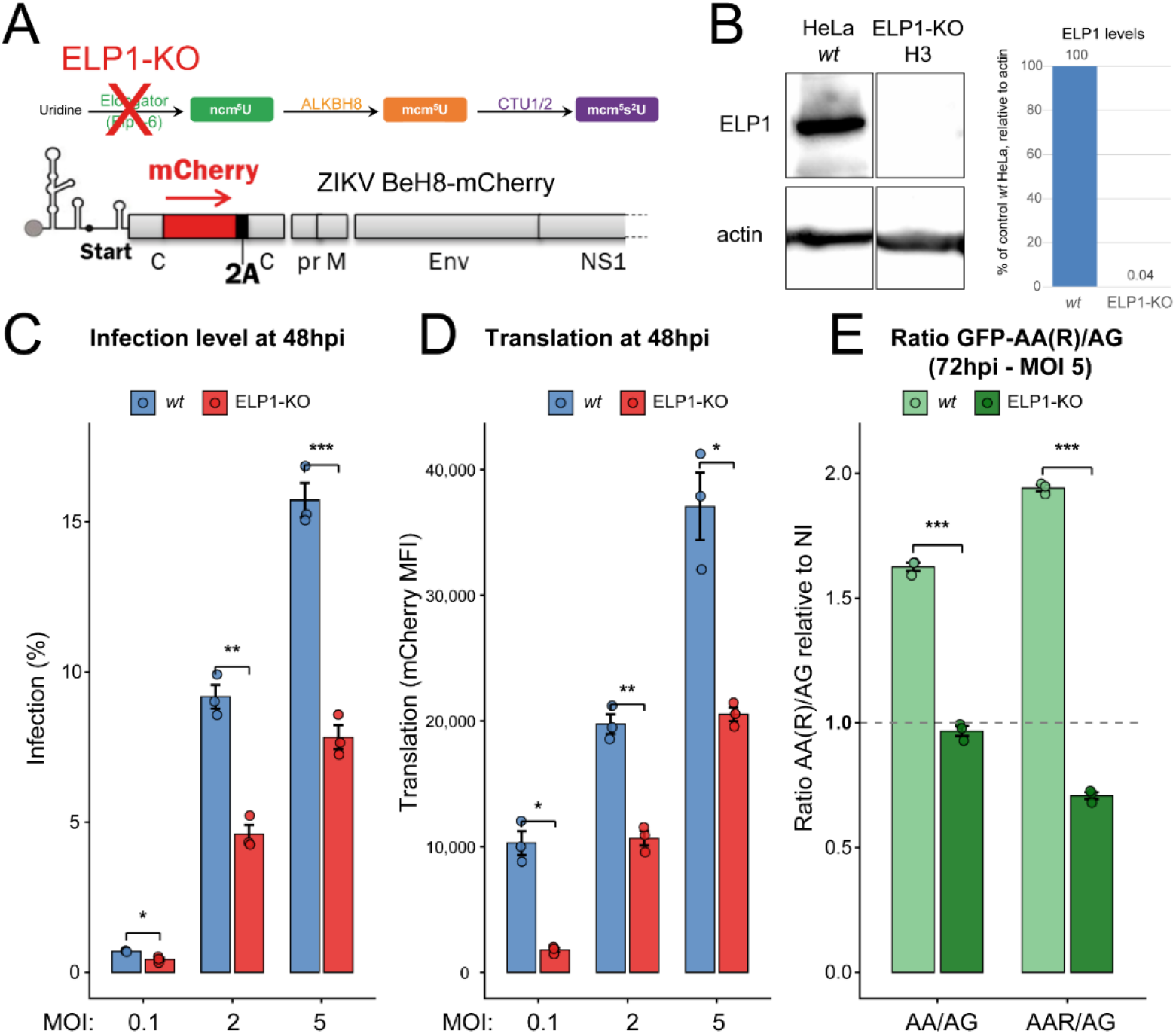
ELP1 knockout impairs U_34_-dependent translation during ZIKV infection. (A) U_34_ pathway is disrupted in ELP1-KO cells leading to the absence of mcm^5^s^2^U modifications on U_34_-containing tRNAs. Partial schematic of ZIKV BeH8-mCherry virus (see *Supplementary Material Figure S2* for construct details). (B) Validation of ELP1-knockout. *Left*, western blot confirming the absence of ELP1 protein in ELP1-KO cells compared to wild-type (wt) HeLa cells. Actin is shown as a loading control. *Right*, the bar graph quantifies ELP1 levels relative to actin, confirming successful knockout. (C) ELP1-KO cells display a significant reduction in infection levels compared to wild-type cells at all MOIs. (D) ELP1-KO cells show a marked decrease in mCherry expression compared to wild-type cells, reflecting impaired viral translation. (E) Codon-dependent translation of GFP sensors in wt and ELP1-KO cells. Ratios of GFP-AA/AG and GFP-AAR/AG translation in wild-type and ELP1-KO cells at 72 hpi (MOI 5), measured using the RNA transfection model with GFP and BFP sensors (as described in Figure 3). Statistical significance: *p < 0.05, **p < 0.01, ***p < 0.001.

Collectively, these findings demonstrate that U_34_ tRNA modifications are essential for optimal ZIKV translation, affecting both infection efficiency and codon-dependent protein synthesis. The impaired translation of codon-biased constructs in ELP1-KO cells, particularly for Arg^*AGA*^-rich sequences, underscores the importance of mcm^5^s^2^U modifications in supporting ZIKV’s codon-specific replication strategy.

## Discussion

Our findings identify a previously unrecognized mechanism by which ZIKV exploits host U_34_ tRNA modifications to overcome its codon usage bias, representing a novel layer of viral adaptation within the expanding field of epitranscriptomic regulation of virus-host interactions. Bioinformatic analysis indicate that ZIKV exhibits a strong preference for A-ending codons, which contrasts with the human host’s preference for G-ending codons, confirming its poor adaptation of codon usage towards its host (70). Since the proper decoding of A-ending codons relies on specific modifications at the wobble position (U_34_) of the cognate tRNAs, we postulated that ZIKV manipulates host tRNA U_34_ modifications to compensate for its non-optimal codon usage, thereby enhancing translation efficiency.

Standard bioinformatic metrics (CAI, RSCU, and Enc) have previously revealed a weak codon bias in Zika virus (ZIKV), with minimal changes in CAI observed during its evolution (71). However, specific studies have highlighted the adaptation of the NS1 protein to human hosts (72), as well as a higher codon usage bias in the NS2B and NS4A proteins (73). These patterns are evident in the CAI profile of ZIKV strain PF13, shown in Figure 2A: the NS1 gene segment exhibits CAI peaks exceeding 0.9, while NS2B and NS4A display CAI values dropping below 0.6. Additionally, previous research has documented distinct codon preferences in ZIKV genes relative to human codon usage, with a notable predominance of A-ending codons among the most frequently used (13). RSCU analysis further revealed codon usage differences between Asian strains and the historical African isolate MR766. While MR766 codon usage remains distinct from the host preferences, its enrichment in U_34_-sensitive codons is less pronounced than that of Asian strains. Notably, MR766 is known to be more virulent, inducing lower innate and inflammatory activation(74) than Asian lineages, leading to more acute infections and more severe brain damages (75, 76). In light of the recent resurgence of MR766-related African strains in West Africa(77, 78), additional investigation is warranted to determine whether codon bias disparities influence the phenotypic characteristics of contemporary African strains(79, 80). The pronounced mismatch between ZIKV codon usage and host tRNA availability suggests a highly specialized evolutionary strategy. Our analysis indicates that ZIKV has evolved a strong preference for A-ending codons—such as Lys^*AAA*^, Glu^*GAA*^, Gln^*CAA*^, and notably Arg^*AGA*^—which are infrequently utilized in the human genome. The contribution of Arg^*AGA*^ codons in the ZIKV genome can be particularly relevant to codon-dependent translation, as demonstrated by our GFP-AAR mRNA sensor experiments. At 72 hours post-infection, the AAR/AG ratio was more enhanced than the AA/AG ratio, indicating that ZIKV infection broadly promotes the decoding of U_34_-sensitive codons, including Arg^*AGA*^. This further supports the idea that ZIKV exploits host tRNA modification pathways to optimize its protein synthesis. Our results from Figure 7 further emphasize the critical role of U_34_ modifications in ZIKV translation. ELP1-knockout cells, which lack mcm^5^s^2^U modifications on U_34_-containing tRNAs, exhibited significantly reduced infection levels, impaired viral translation, and diminished decoding of codon-biased GFP constructs, particularly for Arg^*AGA*^-rich sequences. This underscores the preponderant contribution of Arg^*AGA*^ codons in ZIKV’s codon-specific translation strategy.

The presence of high-order U_34_-sensitive codon stretches (quadruplets and quintuplets) within NS1, NS3, and NS5 suggests that these clusters could function as translational checkpoints. By leveraging the host’s tRNA modification levels, ZIKV may regulate the speed of its replication machinery. Similar to the translational checkpoint identified around the 50th codon of all four Dengue virus (DENV) serotypes(81), this ‘bottleneck’ strategy might prevent the premature overproduction of viral proteins—a hypothesis that remains to be formally tested in future studies.

The use of codon-biased GFP sensors, designed to reflect either human or viral gene preferences, strongly indicated that ZIKV can manipulate the host translation machinery to promote its own protein synthesis. This was supported by analysis of tRNA epitranscriptome dynamics during ZIKV infection, which revealed a transient increase in the major U_34_ tRNA modification, mcm^5^U and mcm^5^s^2^U. Regarding the seemingly modest fold changes (1.1 to 1.2) observed in the total tRNA population, it is important to note that mcm^5^U/mcm^5^s^2^U-modifiable tRNAs constitute only roughly 6–7% of the total tRNA pool (see *Supplementary Figure S10*). Consequently, a 1.1- to 1.2-fold increase across the entire population actually reflects a substantial 2.5- to 4.5-fold enrichment specifically within the U_34_-modifiable tRNA subset. This striking increase suggests that these specific tRNAs may be approaching near-saturation levels of modification.

In our system, ZIKV infection triggers an early and transient increase in wobble mcm^5^U/mcm^5^s^2^U levels followed by a progressive decline at later time points. Although no enzyme has been conclusively shown to directly demethylate wobble mcm^5^U/mcm^5^s^2^U in tRNAs, dynamic changes in this modification are well in line with current models of stress-dependent remodeling of the tRNA modification landscape (82). Multiple studies have demonstrated that cellular stress reprograms U_34_ and other tRNA modifications through regulation of tRNA-modifying enzymes, thereby tuning translation of stress-responsive transcripts rather than relying on dedicated “eraser” activities for each modification (83). We hypothesize that the late decrease in mcm^5^s^2^U we observed can be a consequence of altered tRNA modification flux under ZIKV-induced stress—combining reduced activity or expression of mcm^5^s^2^ writer enzymes (Elongator, ALKBH8, CTU1/CTU2), changes in precursor and cofactor availability, and differential stability or turnover of highly modified tRNAs—rather than as active demethylation of pre-existing mcm^5^s^2^U (84, 85). This view is further supported by work showing that loss or attenuation of mcm^5^s^2^U formation, for example in Trm9/Trm112(83, 86) or sulfur-transfer mutants (87), leads to marked changes in stress signaling and redox homeostasis, underscoring that downmodulation of mcm^5^s^2^U levels can arise from downregulation of the biosynthetic pathway itself(88).

The critical role of this modification was underscored by the severely impaired ZIKV translation observed in ELP1-deficient human fibroblasts, which are intrinsically unable to generate U_34_-modified tRNAs. Remarkably, treatment with RECTAS, a small molecule that restores U_34_ modifications, reversed this phenotype and significantly enhanced ZIKV translation when applied to infected cells. Furthermore, disruption of U_34_-modifying enzymes via CRISPR or shRNA approaches consistently impaired ZIKV replication, underscoring the critical requirement of the mcm^5^s^2^U modification machinery in viral translation. Building on the mechanistic insight provided by RECTAS, which promotes exon 20 inclusion in *ELP1* transcripts of FD cells(44, 89), a new therapeutic strategy involving the identification of derivative molecules capable of excluding exon 20 in wild-type ELP1 mRNA in normal cells can be conceptualized, as we recently exemplified in the context of SARS-CoV-2 infection through splice-switching of OAS1 isoforms(90). Exclusion of ELP1 exon 20 could reduce U_34_ tRNA modifications, thereby disrupting ZIKV translation and potentially curbing infection. However, a significant challenge remains in achieving antiviral efficacy without inducing host cell toxicity, as this strategy targets a mechanism used by both host and virus.

Viruses are highly dependent on the host cell’s translation machinery, including host tRNAs, for efficient protein synthesis(4, 10, 91). Some viruses can alter tRNA abundance as part of their adaptive strategy (92), as described by us for SARS-CoV-2(46) and by others for retroviruses such as HIV(93) and DNA viruses, including SV40(94), EBV(95), Adenovirus(96), and HSV-1(97). In addition, viral translation is influenced by chemical modifications of tRNAs, particularly within the anticodon loop region, catalyzed by a suite of tRNA-modifying enzymes(36, 98). Notably, we have already shown the ability of SARS-CoV-2 to directly alter the host tRNA epitranscriptome(99, 100) by upregulating U_34_-modified tRNAs that can optimize the translation of its genome, which is enriched in U_34_-sensitive codons(46). This mechanism, whereby viruses reprogram tRNA modifications to favor the decoding of their codon-biased genomes, has since been independently confirmed(101), showing that SARS-CoV-2 infections induce stress-responsive alterations including a net increase of tRNA mcm^5^U/mcm^5^s^2^U levels, thereby enhancing the translation of viral proteins enriched in A-ending codons. A similar strategy has been proposed for other RNA viruses, including Chikungunya virus (CHIKV) and Dengue virus (DENV), though direct experimental validation remains limited(14, 102–104). The transient increase of tRNA mcm^5^U/mcm^5^s^2^U levels was also reported during influenza A virus infections (105). Notably, all observed variations in mcm^5^U/mcm^5^s^2^U levels were comparable in magnitude to those detected here during ZIKV infection. Our findings now extend this concept to ZIKV, and our functional analyses provide some of the most direct evidence to date of this mechanism in action, reinforcing the idea that tRNA epitranscriptome reprogramming is a conserved strategy among RNA viruses to optimize viral protein synthesis.

While a universal mechanism used by different RNA viruses to exploit U_34_ modifications for their benefit has not been identified, recent studies have demonstrated that infection by the Alphavirus CHIKV upregulates the expression of KIAA1456, an ALKBH8 homolog responsible for generating the mcm^5^U_34_ modification on tRNAs. This enhancement results in reprogramming codon optimality and favors CHIKV RNA translation(102). DENV exploits host tRNA through a different mechanism by manipulating the levels of ALKBH1, which introduces 2’-O-methyl-5-formylcytidine (f^5^Cm) modification at the wobble position C_34_ of cytoplasmic tRNA-Leu(CAA), promoting viral translational remodeling during DENV infection(103). Alterations in the tRNA epitranscriptome induced by Coronaviruses (SARS-CoV-2, HCoV-OC43) notably including mcm^5^s^2^U upregulation (101) are potentially linked to host stress pathways, such as oxidative stress and DNA damage responses (DDR), which activate key signaling molecules (e.g., phosphorylated ATM, γH2AX, and p38 MAPK) and ultimately promote expression of U_34_ enzymes (ELP3, KIAA1456 or its paralog ALKBH8) leading to an enhanced decoding of A-ending codons enriched in Coronavirus genomes. Given the ability of ZIKV infection to induce Endoplasmic Reticulum (ER) stress(106), Unfolded Protein Response (UPR)(107), and oxidative stress(108, 109), it is plausible that ZIKV exploits similar mechanisms by hijacking host stress pathways to enhance mcm^5^s^2^U_34_ levels and optimize viral translation. The PI3K-Akt-mTORC2 signaling pathway, known to reprogram the translation of genes enriched in U_34_-sensitive codons by phosphorylating the ELP1 subunit of the Elongator complex at Ser1174(110, 111), may also play a role in this process. ZIKV NS4A/NS4B viral proteins have been implicated in deregulating the Akt-mTOR pathway(112), suggesting a potential molecular connection to tRNA reprogramming during infection. Furthermore, the virus-induced oxidative stress and ER stress could synergistically enhance mcm^5^s^2^U_34_ modifications, similar to the mechanism described for SARS-CoV-2(101). This finding provides a compelling avenue for future investigations, particularly in exploring whether ZIKV infection triggers ATM/ATR activation, ROS production, or other stress pathways that could drive tRNA epitranscriptome remodeling.

Finally, whether ZIKV-mediated U_34_ alterations impact the host cell metabolism is also an open question. ZIKV infection is strongly associated with microcephaly in infants born to infected mothers, as well as Guillain-Barré Syndrome (GBS) in adults, cognitive disability, and symptoms of autonomic dysfunction(113). In this study we focused on brain-resident cells such as astrocytes, the most abundant neuronal cells targeted by ZIKV(114). The ZIKV-driven enhancement of the translation of genes enriched in A-ending codons at the expense of genes preferring G-ending codons, may profoundly alter the cellular proteome integrity(115). Indeed, among disease-relevant modifications affecting cytoplasmic tRNA species, neurological disorders including Familial Dysautonomia (FD)(68, 116) and epilepsy, appear to be most common. Specifically, mutations impairing m^5^C, mcm^5^U, mcm^5^s^2^U, ncm^5^U, pseudouridine (Ψ), m^1^G, inosine (I)(117) and m^2^_2_G(99, 118) tRNA modifications are all associated with intellectual disability and neurodevelopmental disorders, suggesting normal brain function and development to be particularly dependent on the presence of these modifications(18). In many aspects, the neurological consequences of ZIKV infection, that are still barely understood, might be prospectively phenocopying (to a certain extent) alterations related to U_34_ enzymes and tRNA modifications.

In conclusion, our findings place ZIKV within a growing group of RNA viruses that modulate host tRNA modifications as a strategy to facilitate viral protein synthesis. The virus’s reliance on the mcm5U/mcm^5^s^2^U tRNA modification pathway highlights a critical vulnerability that may be therapeutically exploitable, albeit with caution due to the essential nature of these modifications in host physiology.

## Acknowledgements

This work was initially facilitated by a Hubert Curien exchange program from Campus France (PHC Aurora 2018 - EpiArbo N°40969YB) between PE and GS. PROMEC is a member of the National Network of Advanced Proteomics Infrastructure (NAPI), which is funded by the RCN INFRASTRUKTUR-program (295910).

## Supplementary Data

are available at NAR Online

The data underlying this article are available in the article and in its online **supplementary material**.

## Author Contributions statement

PE conceived the project, realized the experiments, analyzed the data, generated the figures and wrote the paper. EB technically supported RNA extractions and virus productions. CBV performed mass spectrometry analysis at PROMEC facility. SS provided human primary astrocytes. LG supported the use of codon biased GFP reporters and Familial Dysautonomia target cells. MA and MH provided RECTAS molecules to revert ELP1-deficiency in FD cells. GS analyzed mass spectrometry data and contributed to paper editing. LB participated to the project conception and financial support, analyzed the data and contributed to paper writing and editing.

