## Supplementary figures and images for "Zika virus reprograms the host tRNA epitranscriptome to adapt translation to A-ending codon bias"

### Supplementary data

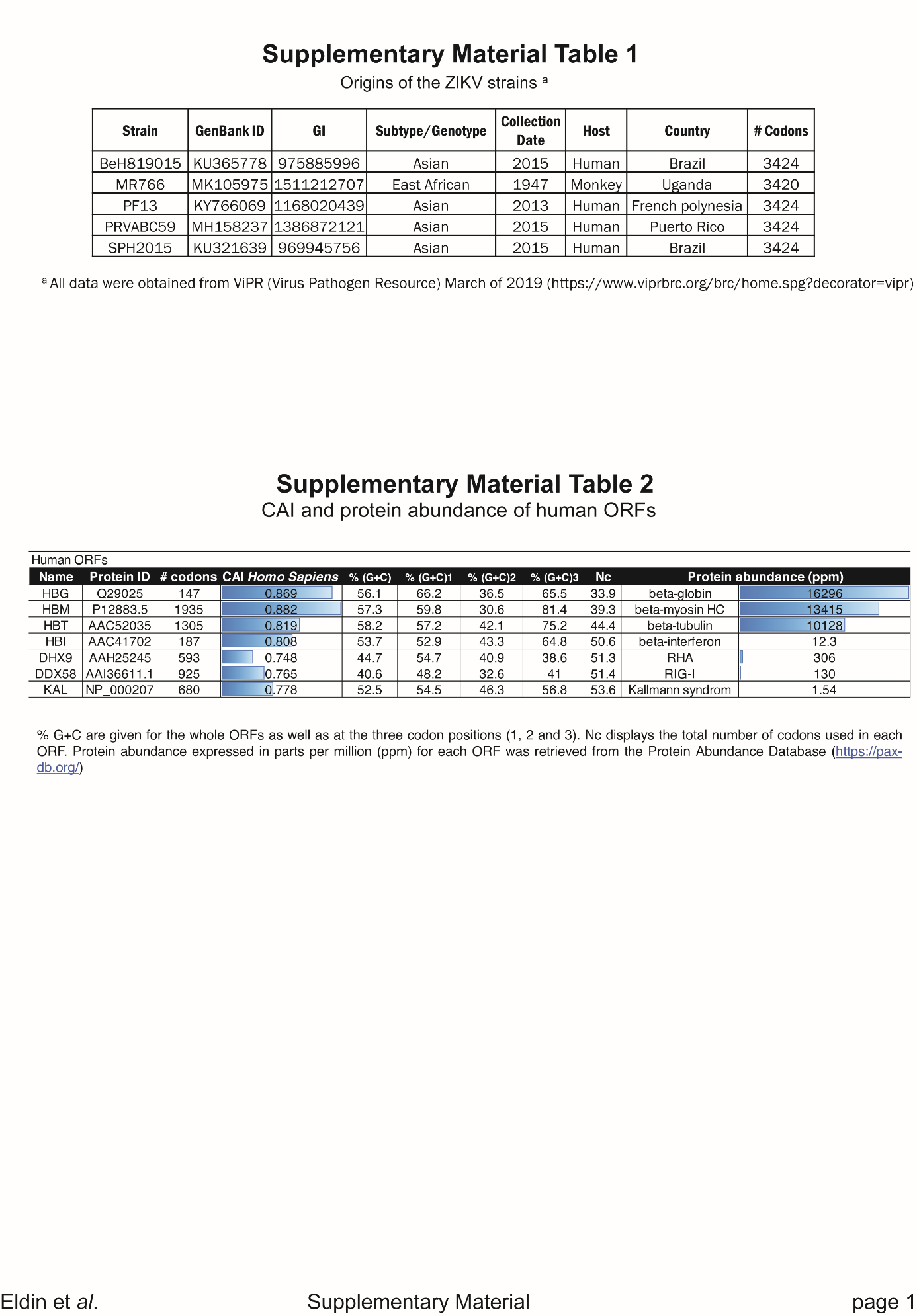

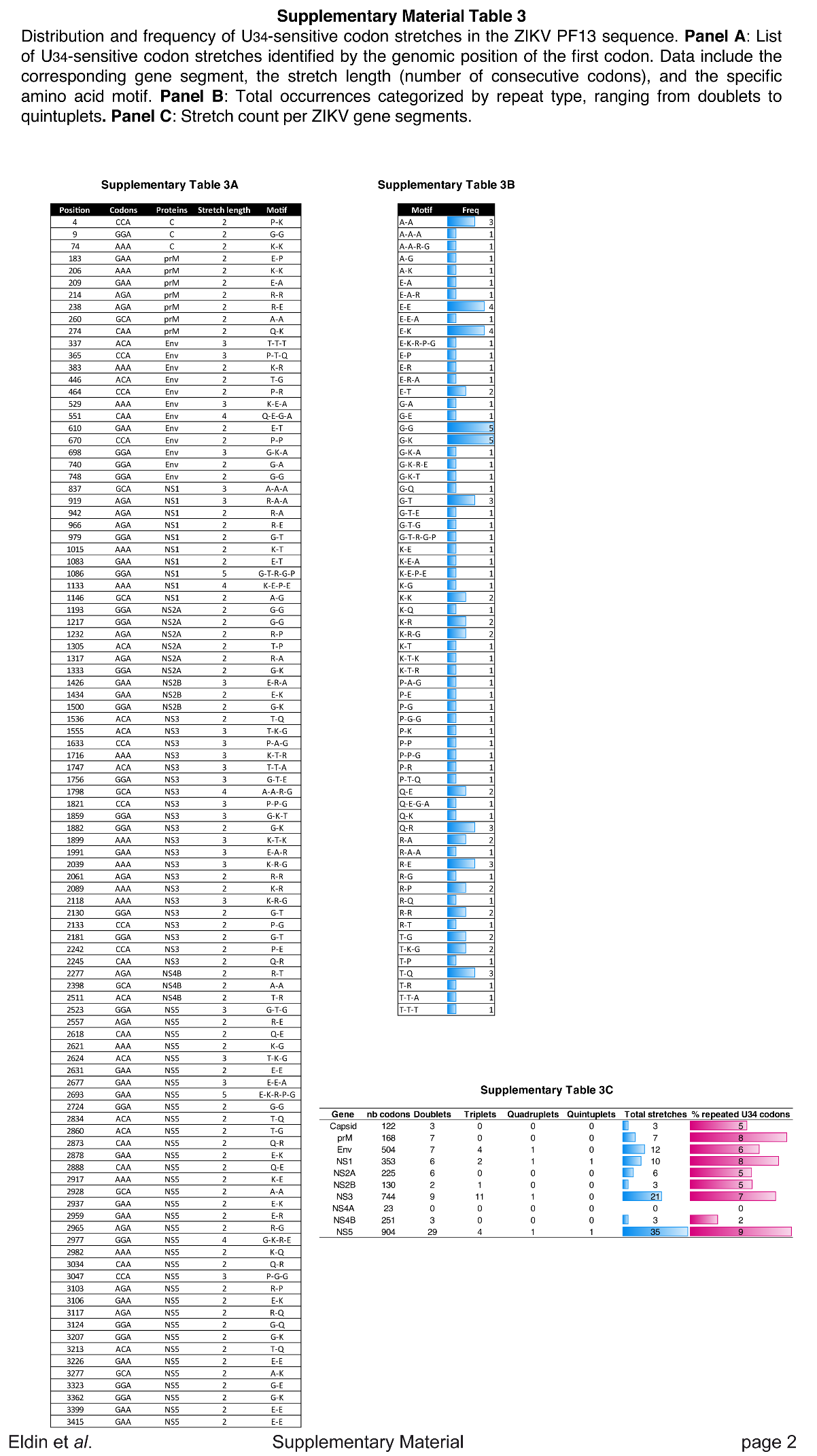

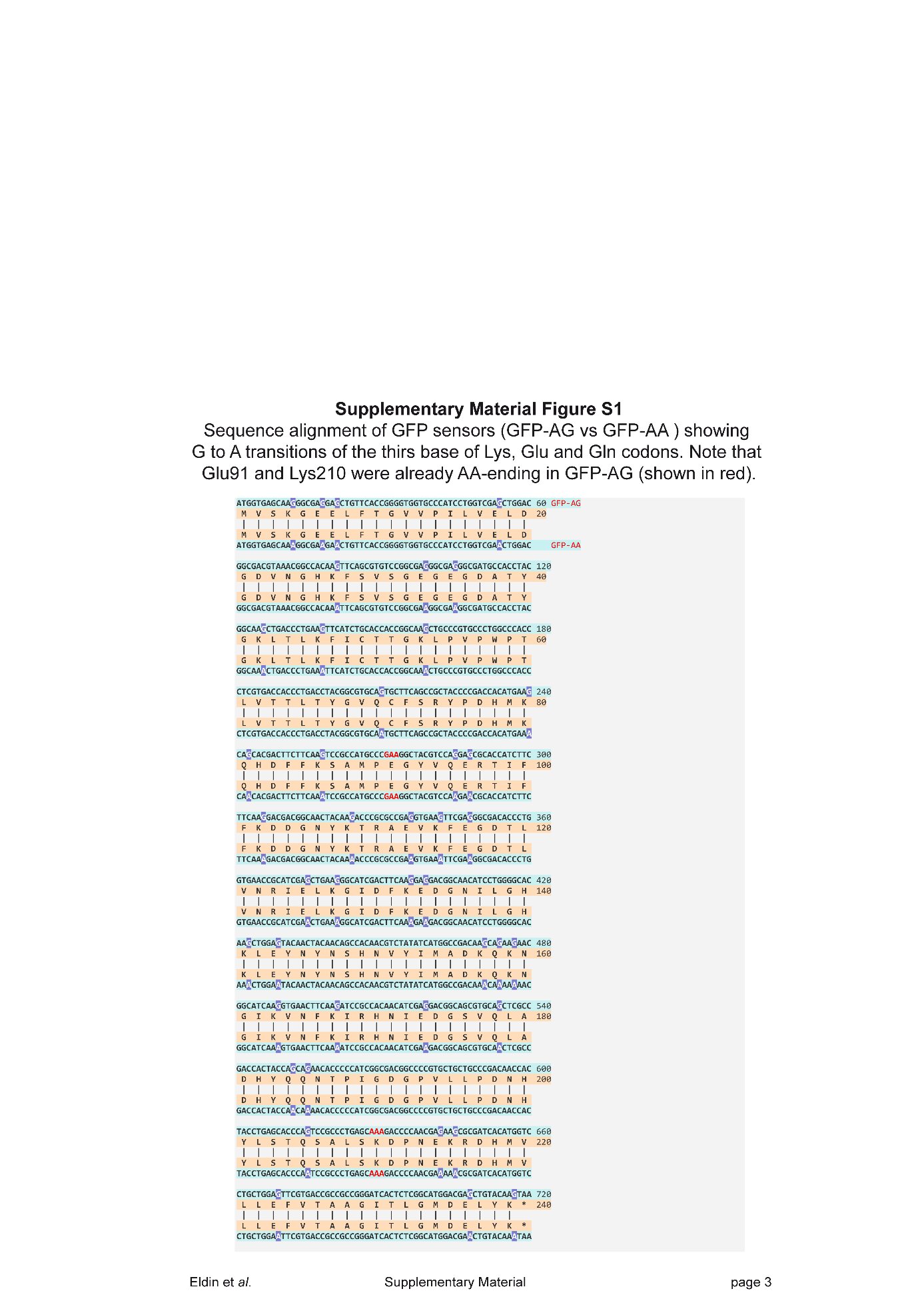

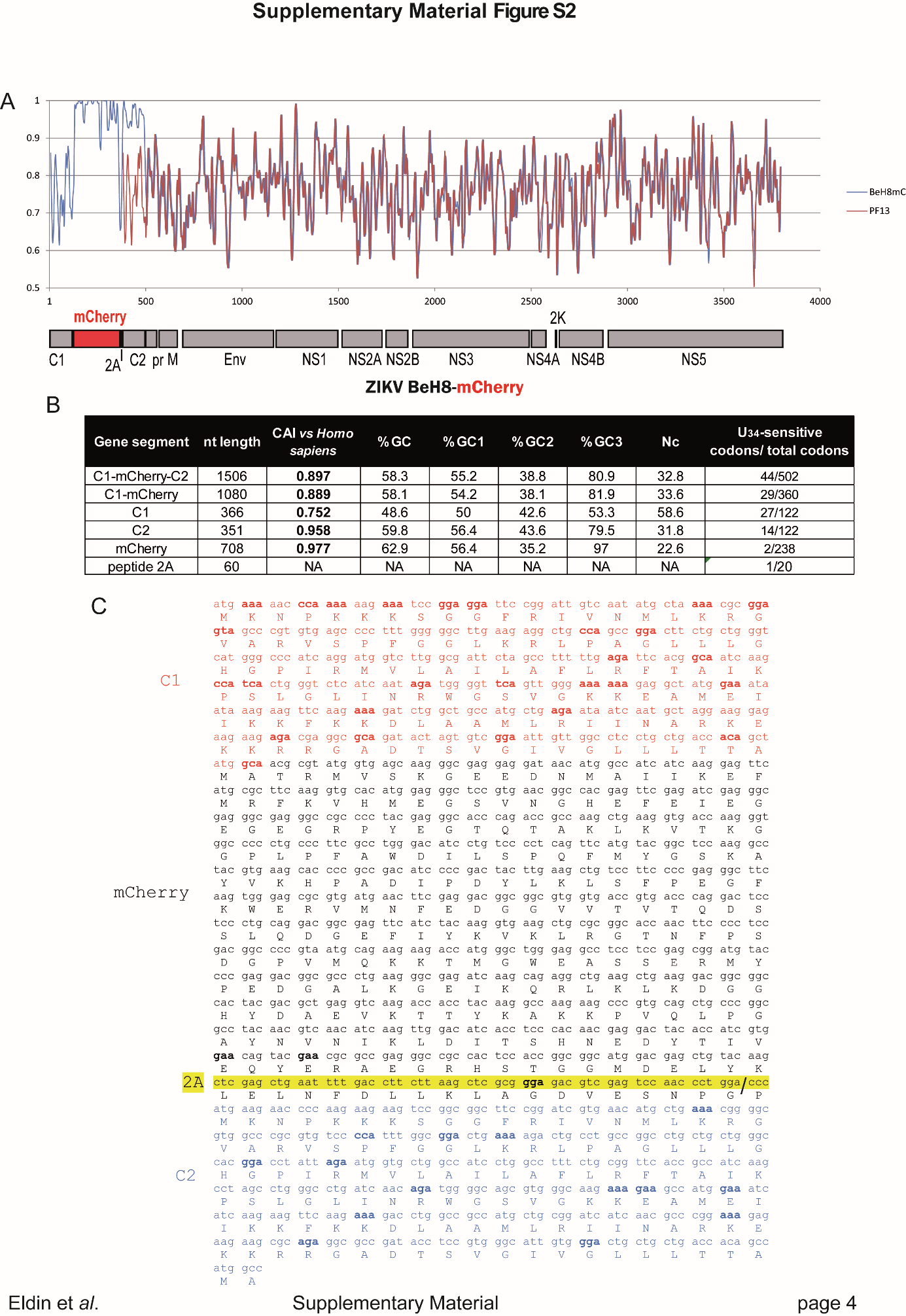

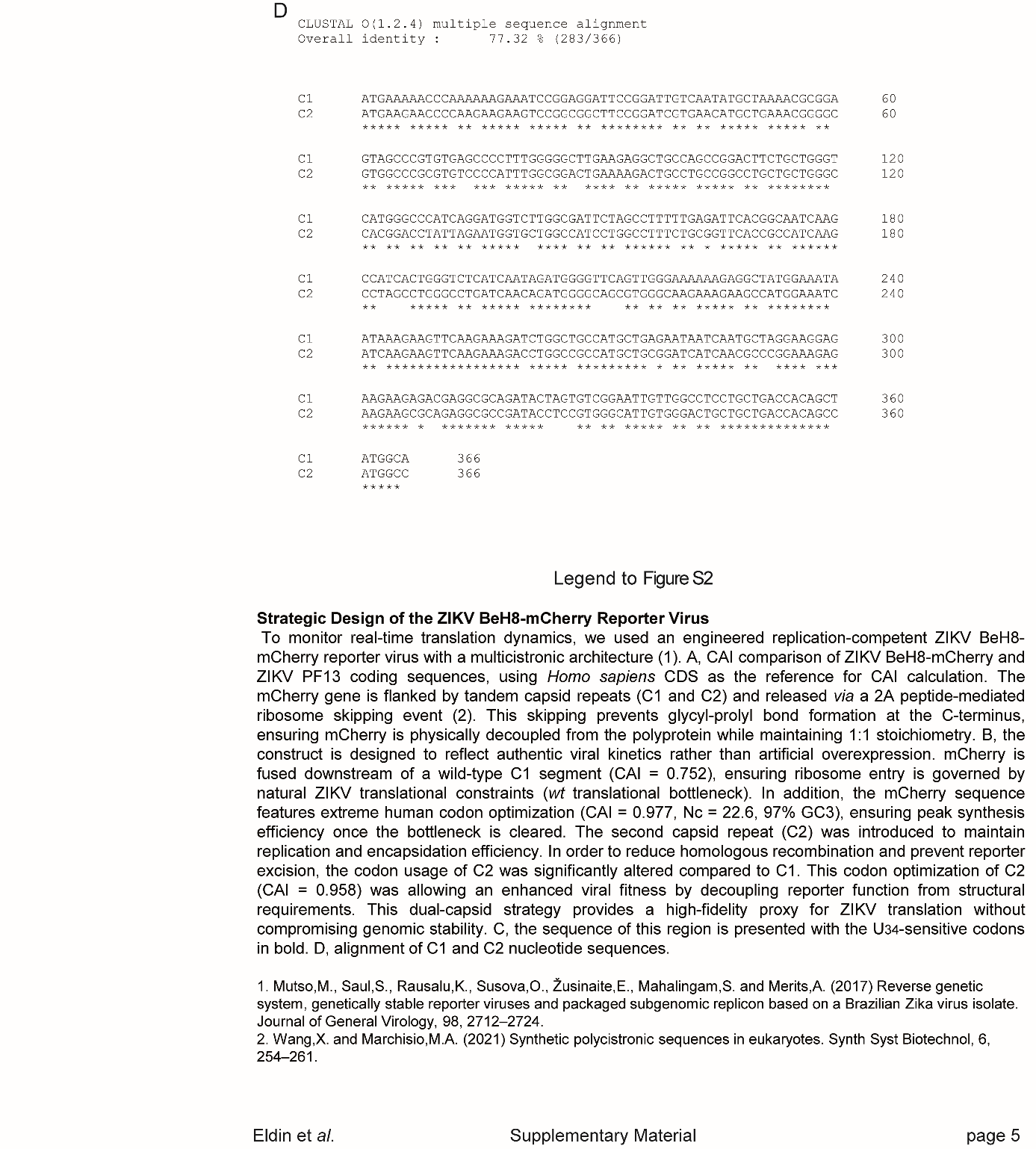

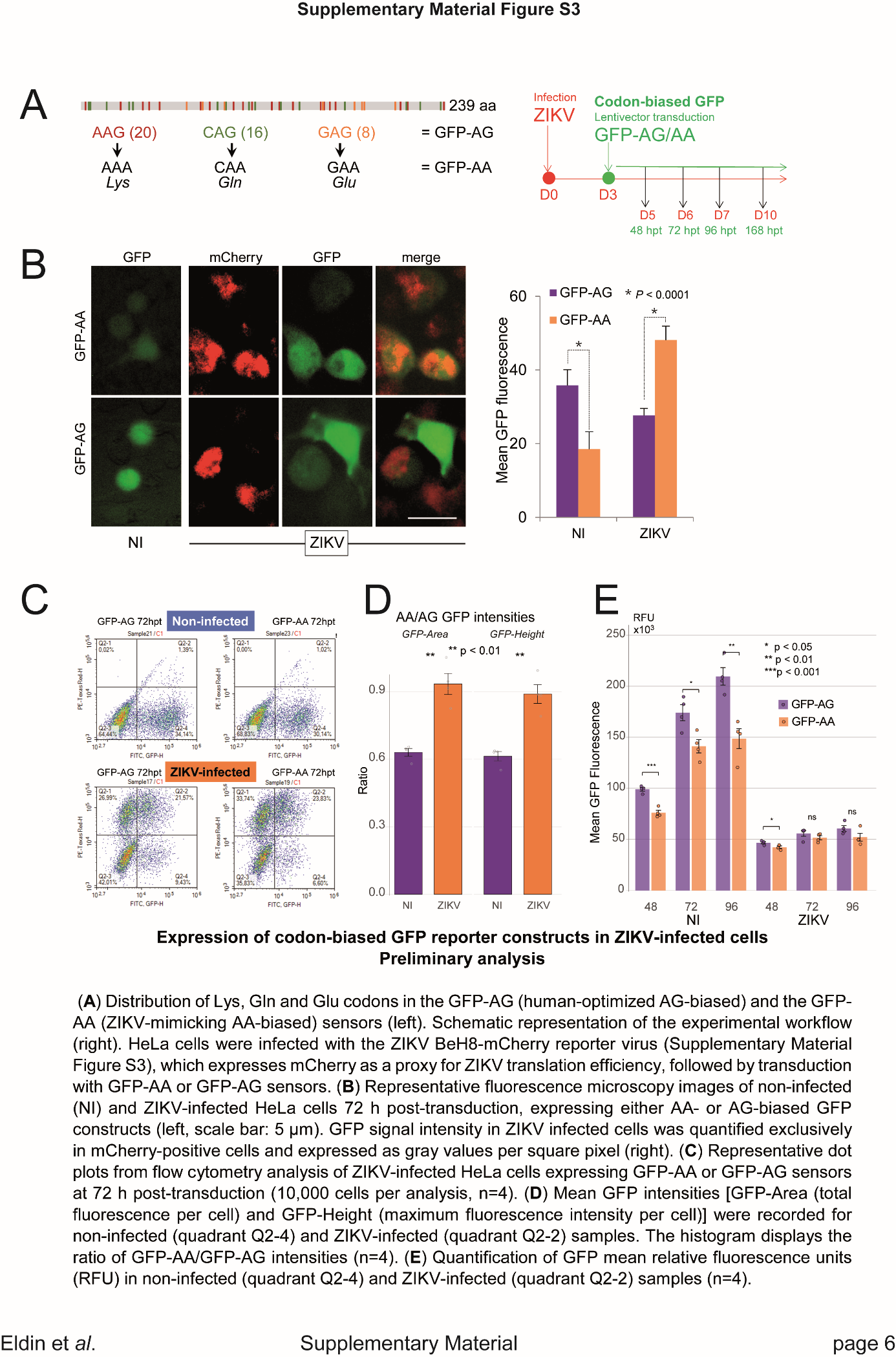

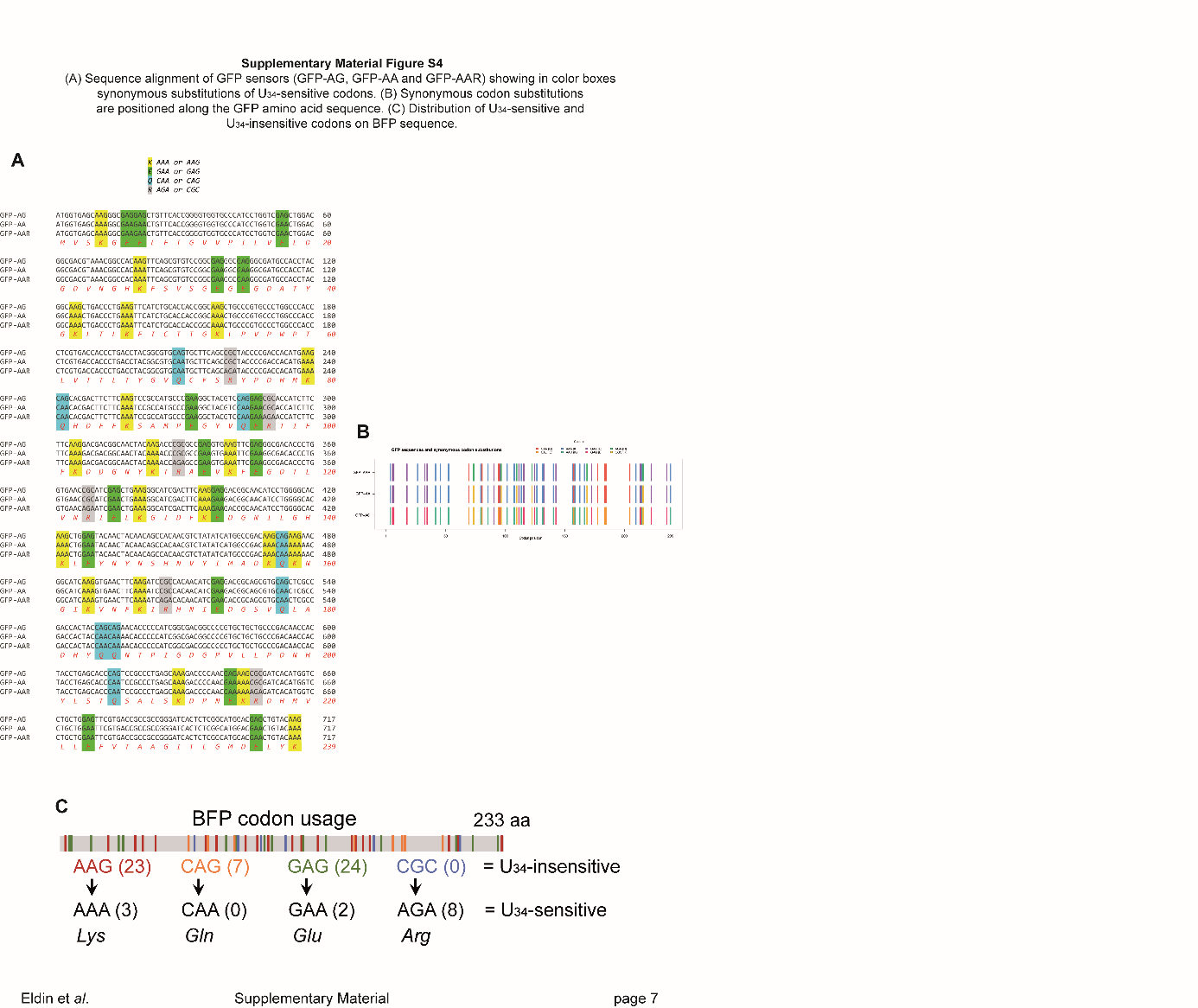

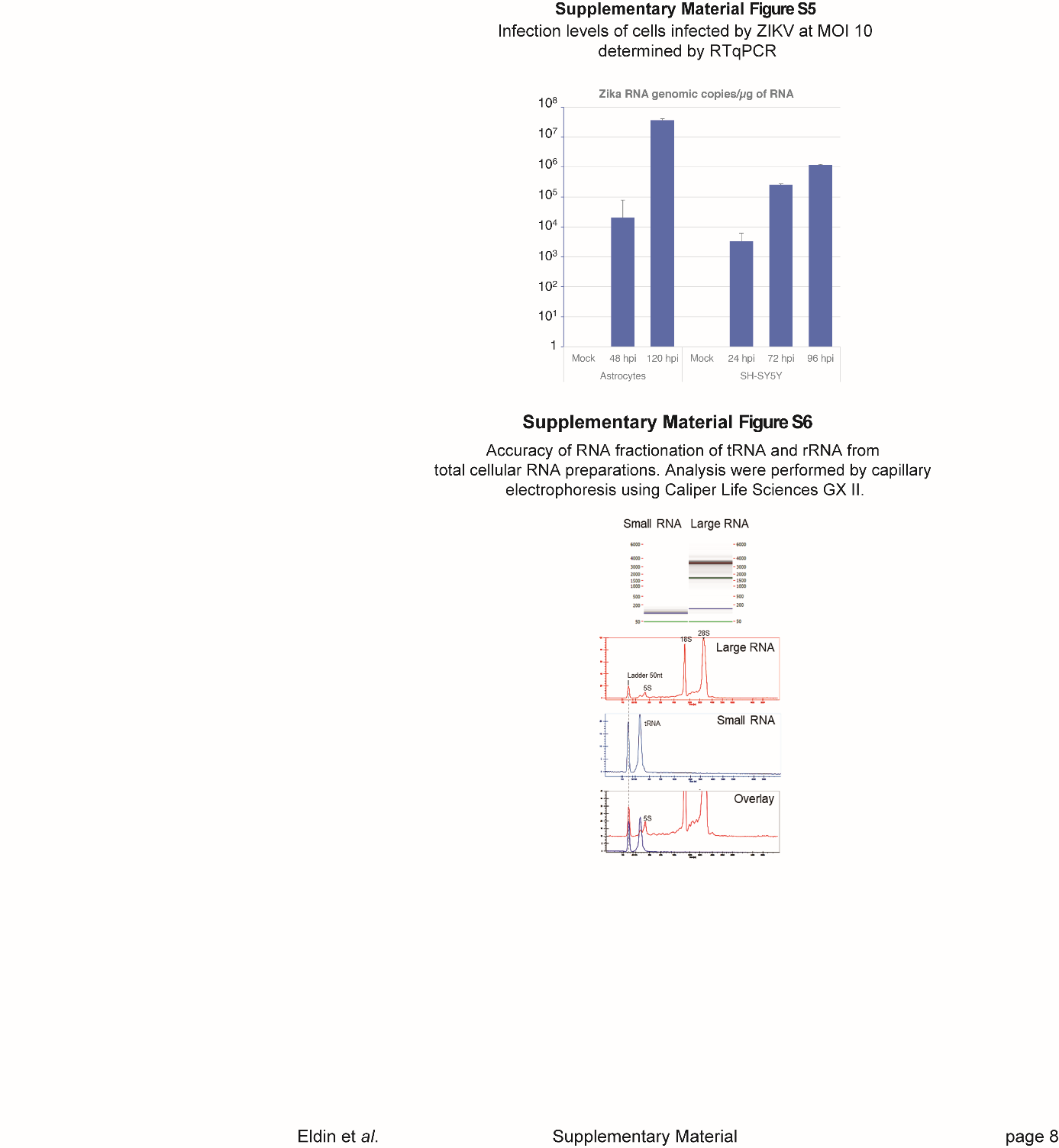

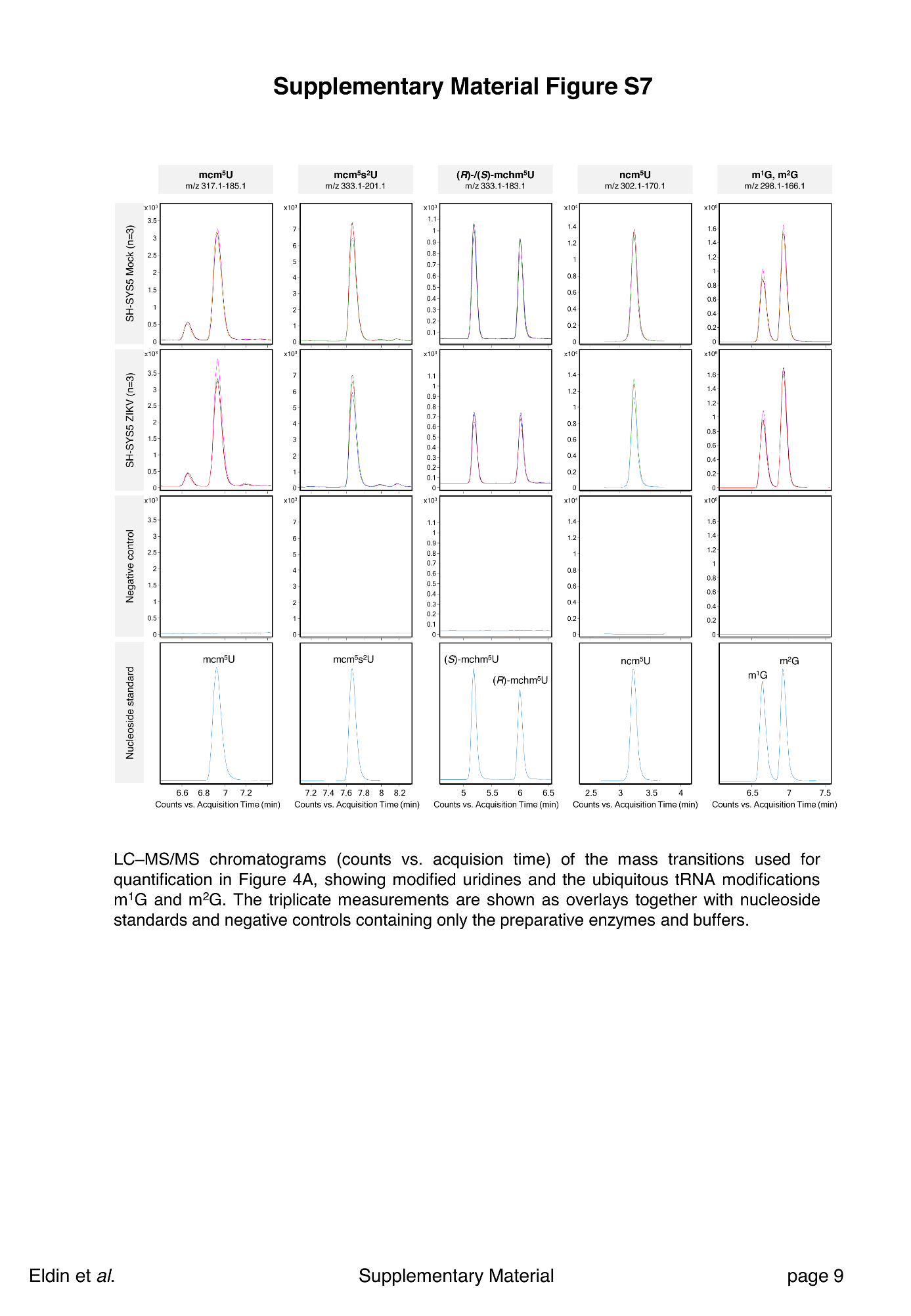

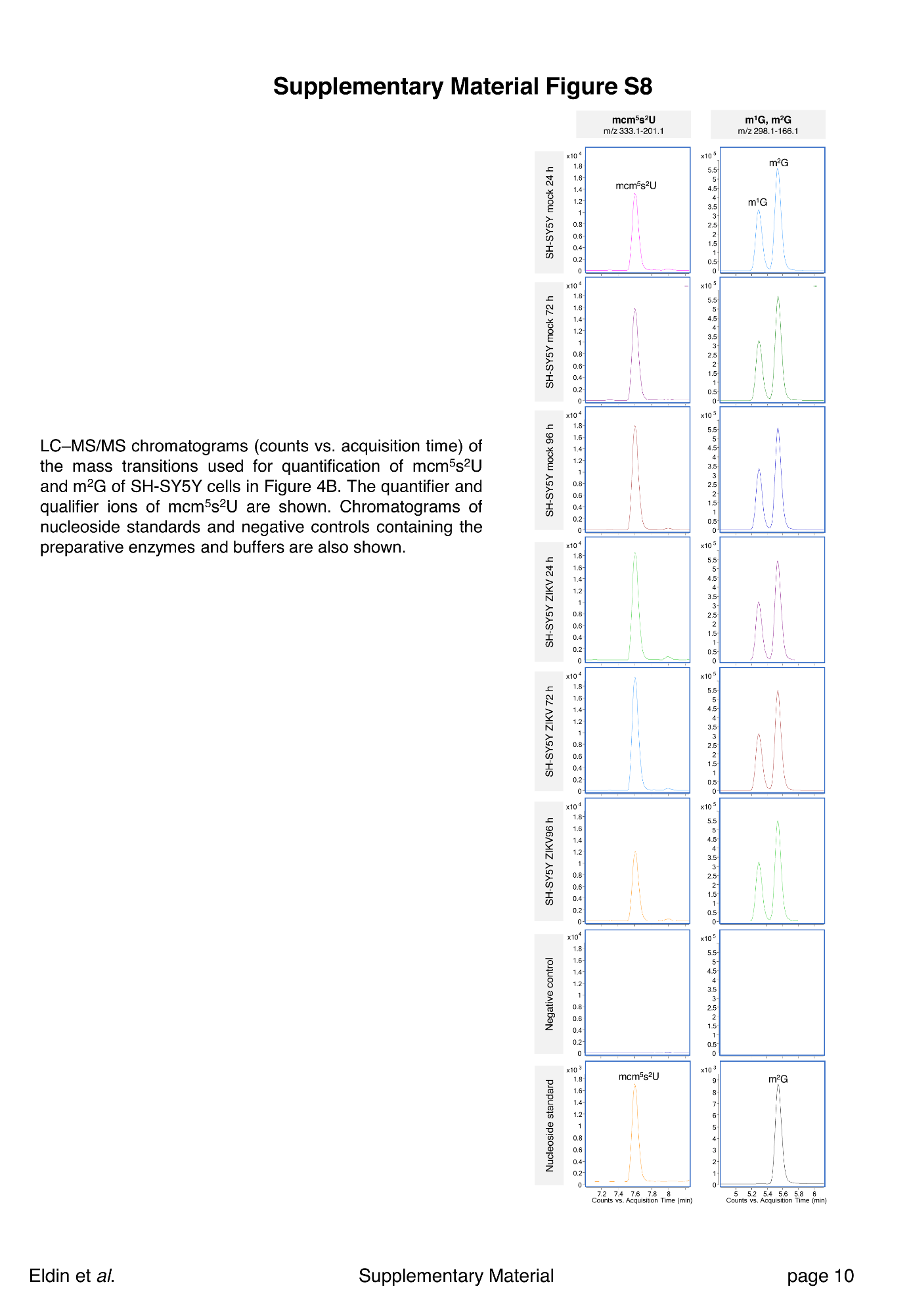

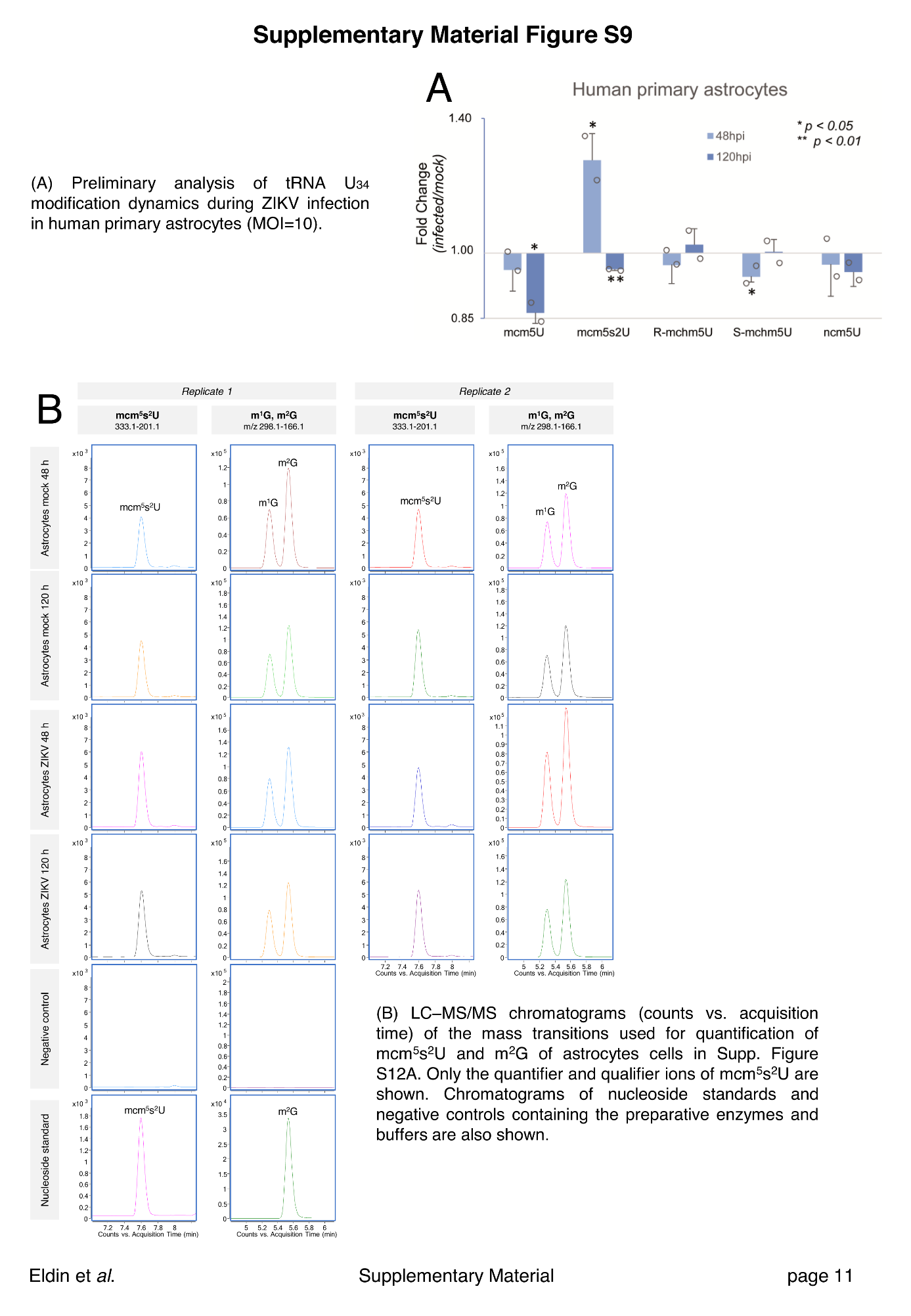

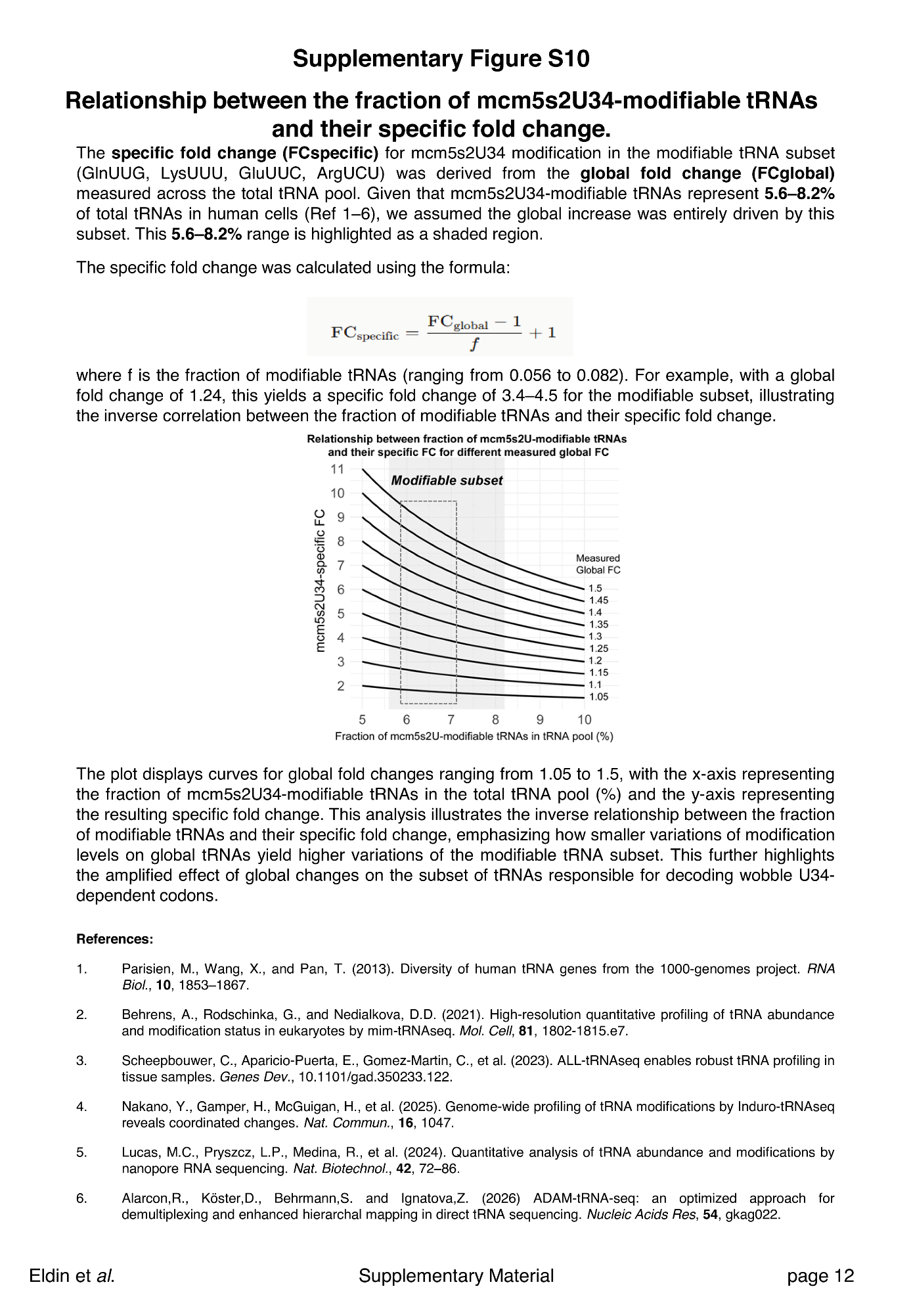

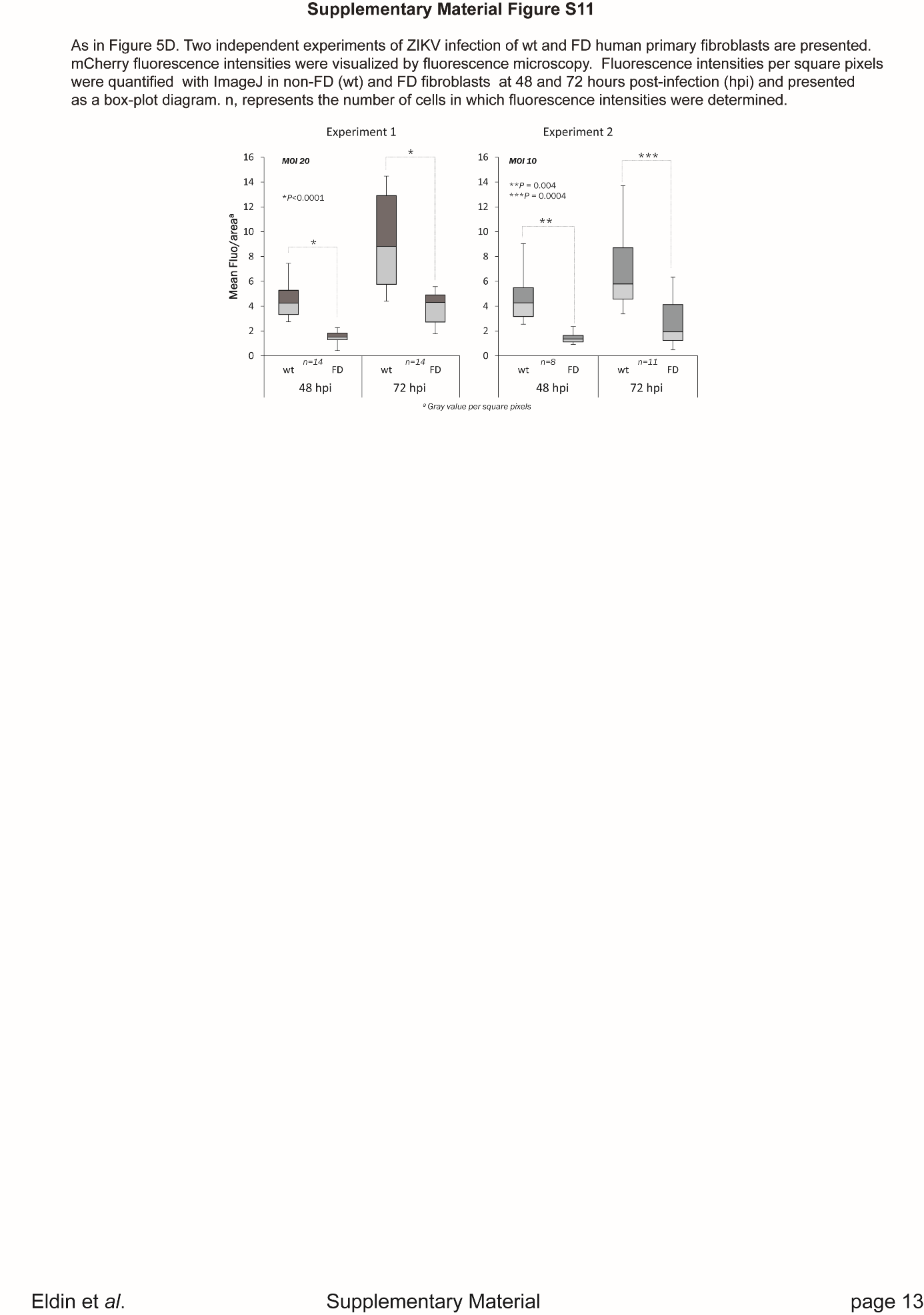

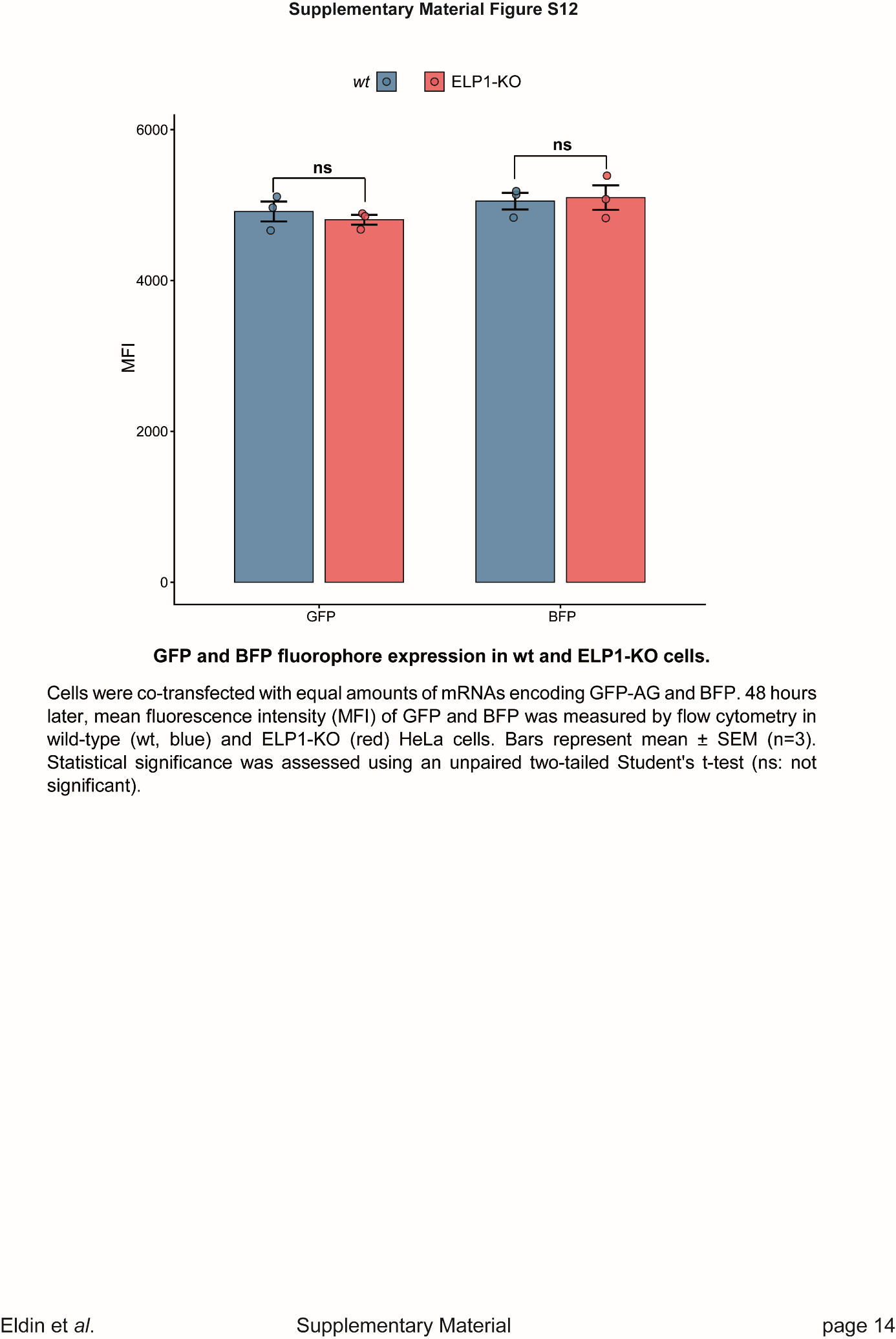

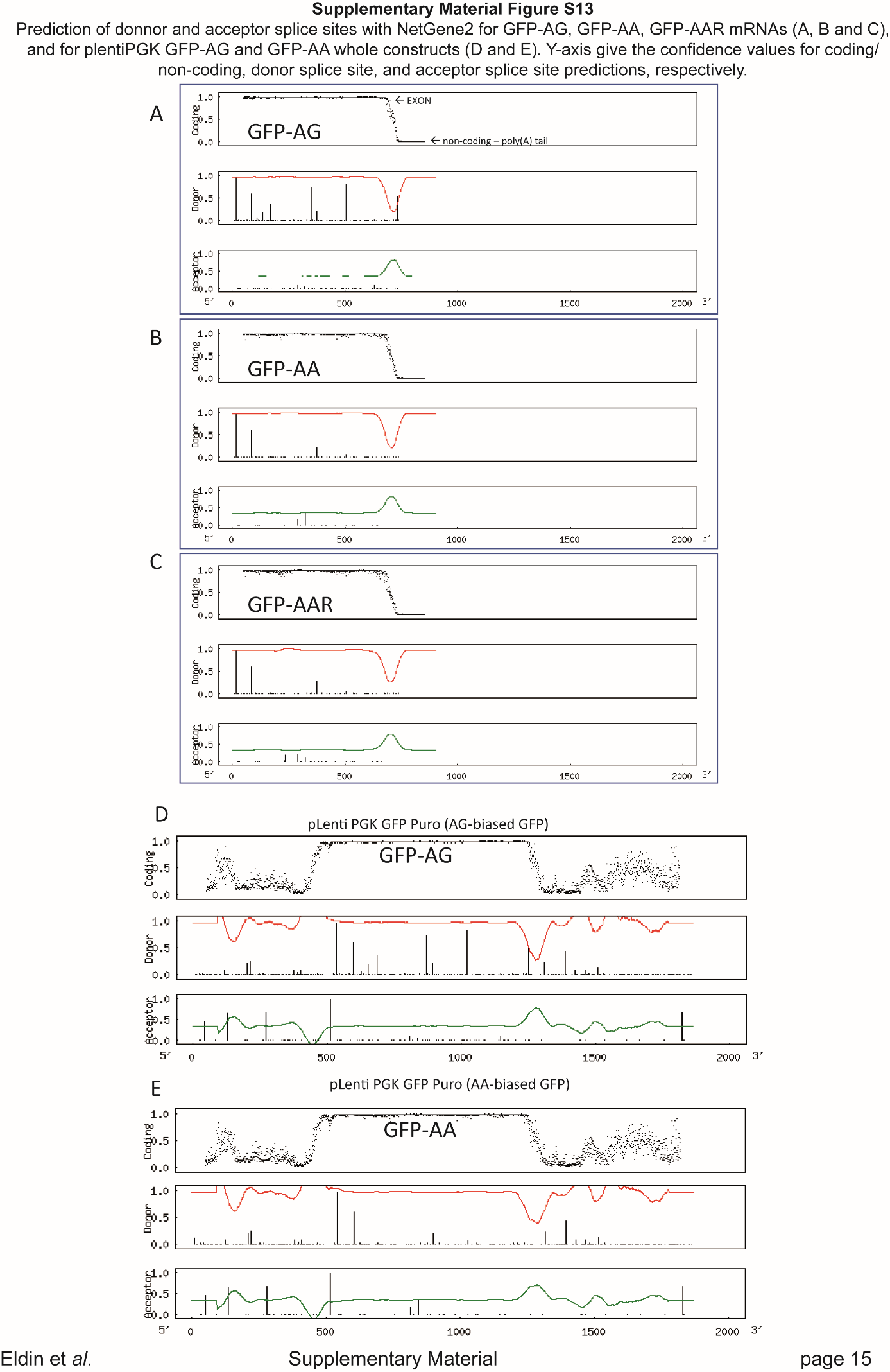
